# Convergent evolution in toxin detection and resistance provides evidence for conserved bacterial-fungal interactions

**DOI:** 10.1101/2023.11.27.568923

**Authors:** Stephen K. Dolan, Ashley T. Duong, Marvin Whiteley

**Affiliations:** School of Biological Sciences, Georgia Institute of Technology, Atlanta, Georgia, USA, Emory-Children’s Cystic Fibrosis Center, Atlanta, Georgia, USA, Center for Microbial Dynamics and Infection, Georgia Institute of Technology, Atlanta, Georgia, USA; Eukaryotic Pathogens Innovation Center, Department of Genetics and Biochemistry, Clemson University, Clemson, South Carolina, USA

**Keywords:** *Pseudomonas aeruginosa*, *Aspergillus fumigatus*, *zinc*, cystic fibrosis, gliotoxin

## Abstract

Microbes rarely exist in isolation, and instead form complex polymicrobial communities. As a result, microbes have developed intricate offensive and defensive strategies that enhance their fitness in these complex communities. Thus, identifying and understanding the molecular mechanisms controlling polymicrobial interactions is critical for understanding the function of microbial communities. In this study, we show that the Gram-negative opportunistic human pathogen *Pseudomonas aeruginosa*, which frequently causes infection alongside a plethora of other microbes including fungi, encodes a genetic network which can detect, and defend against gliotoxin, a potent, disulfide-containing antimicrobial produced by the ubiquitous filamentous fungus *Aspergillus fumigatus*. We show that gliotoxin exposure disrupts *P. aeruginosa* zinc homeostasis, leading to transcriptional activation of a gene encoding a previously uncharacterized dithiol oxidase (DnoP), which detoxifies gliotoxin and structurally related toxins. While the enzymatic activity of DnoP is identical to that used by *A. fumigatus* to protect itself against gliotoxin, DnoP shares little homology to the *A. fumigatus* gliotoxin resistance protein. Thus, DnoP and its transcriptional induction by low zinc represent an example of both convergent evolution of toxin defense and environmental cue sensing across kingdoms. Collectively, these data support disulfide-containing natural products as mediators of inter-kingdom interactions and provide evidence that *P. aeruginosa* has evolved to survive exposure to these molecules in the natural environment.

**Significance statement:** Bacteria and fungi frequently exist as complex, polymicrobial communities during infection. Reconstructing ecological structure in the laboratory is challenging and, consequently, the precise molecular mechanisms which underpin microbial interactions remain elusive. Using a pre-clinical model that mimics the cystic fibrosis lung, we discovered that the bacterium *Pseudomonas aeruginosa* detects and defends against a disulfide-containing toxin produced by the fungus *Aspergillus fumigatus*. In an example of both convergent evolution of toxin defense and environmental cue sensing across kingdoms, we discovered that these organisms use the same cue to produce/sense this toxin, and the same enzymatic mechanism to protect against toxicity. This discovery of convergent evolution provides strong evidence for *P. aeruginosa* exposure to microbially-produced disulfide-containing toxins in natural environments.

---

Many bacterial infections are not the result of colonization by a single microbe, but instead involve polymicrobial communities. Within these communities, co-infecting microorganisms can interact synergistically and/or antagonistically to induce virulence traits, alter the infected niche, increase tolerance to therapeutics, or modulate the host immune response (reviewed in (1–3)). Consequently, insight into the mechanisms used by pathogens to interact is key to developing specific agents or strategies to limit infection.

Interactions between filamentous fungi and bacteria are widely studied in the context of human infection, but the precise molecular mechanisms underlying these interactions are understudied. The airways of patients with cystic fibrosis (CF) are inhabited by a functionally diverse polymicrobial community with vast potential for interspecies interactions (1). *Aspergillus fumigatus* and *Pseudomonas aeruginosa* are two of the most prevalent fungal and bacterial pathogens isolated from the CF airway. *A. fumigatus* and *P. aeruginosa* can co-infect the lungs of people with CF, with co-infection ranging from 2-45% dependent on the patient cohort (4, 5). In addition, there is correlational clinical evidence that co-infection worsens patient outcomes (4, 6–8), and it has been proposed that interactions between *A. fumigatu*s and *P. aeruginosa* are key to this enhanced pathophysiology. Consequently, there have been numerous in vitro studies examining interactions between *A. fumigatu*s-*P. aeruginosa*, revealing that many of the common *P. aeruginosa* metabolites critical for in vitro interactions with other microbes also impact *A. fumigatus* fitness and physiology during in vitro co-culture ((9–21). However, much less is known regarding the impact of *A. fumigatus* on *P. aeruginosa* physiology, although recent evidence indicates that *A. fumigatus* soluble factors can alter *P. aeruginosa* physiology (22–25).

Here, we used the CF respiratory environment as a model to unravel mechanistic features of *A. fumigatu*s-*P. aeruginosa* interactions, specifically how *P. aeruginosa* physiology is altered by the presence of *A. fumigatus*. Comparative transcriptomics during growth in a pre-clinical model mimicking expectorated human CF sputum (26–29) revealed that upon exposure to *A. fumigatus*, *P. aeruginosa* activates a previously uncharacterized gene, PA4170, to levels observed in human CF infection samples. Comparative metabolomics revealed that PA4170 induction during co-culture is mediated by zinc starvation, a result of *A. fumigatus* secretion of the natural product gliotoxin. Functional characterization of PA4170 revealed that this gene encodes a disulfide natural product oxidase, which protects *P. aeruginosa* from the redox-cycling effects of gliotoxin. This is the same mechanism which *A. fumigatus* uses to protect itself from self-intoxication with gliotoxin. These data provide evidence of convergent evolution of toxin resistance in *A. fumigatus* and *P. aeruginosa* and supports disulfide-containing natural products as key mediators of inter-domain microbial interactions in natural environments.

## Results

### *P. aeruginosa* transcriptional response to *A. fumigatus*

*A. fumigatus*-*P. aeruginosa* mixed species biofilms form readily in vitro when *A. fumigatus* is inoculated first and allowed to form hyphae, followed by *P. aeruginosa* addition (22, 30). Building on this existing model, we cultured *A. fumigatus* (1×10^5^ conidia/ml) statically in synthetic cystic fibrosis medium (SCFM2); a defined medium designed to mimic the chemistry and viscosity of expectorated human CF sputum, which promotes *P. aeruginosa* biofilm aggregate formation (27, 29). After 12-16 hr, filamentous growth of *A. fumigatus* was observed in SCFM2, at which point, *P. aeruginosa* (5 × 10^5^ cells mL^−1^) was added to the culture. The co-culture was allowed to grow until *P. aeruginosa* reached early stationary phase (∼12 hours, **Fig. S1A**), at which time it was harvested for transcriptome analysis. No significant growth alterations were noted for *P. aeruginosa* or *A. fumigatus* in coculture when compared to monoculture (**Fig. S1A, S1B**), and there was close association between these microbes during co-culture, as shown previously (13, 31, 32) (**Fig. 1A**). This experimental setup allowed us to examine the transcriptional response of *P. aeruginosa* to co-culture with *A. fumigatus*.

**Fig. 1.**
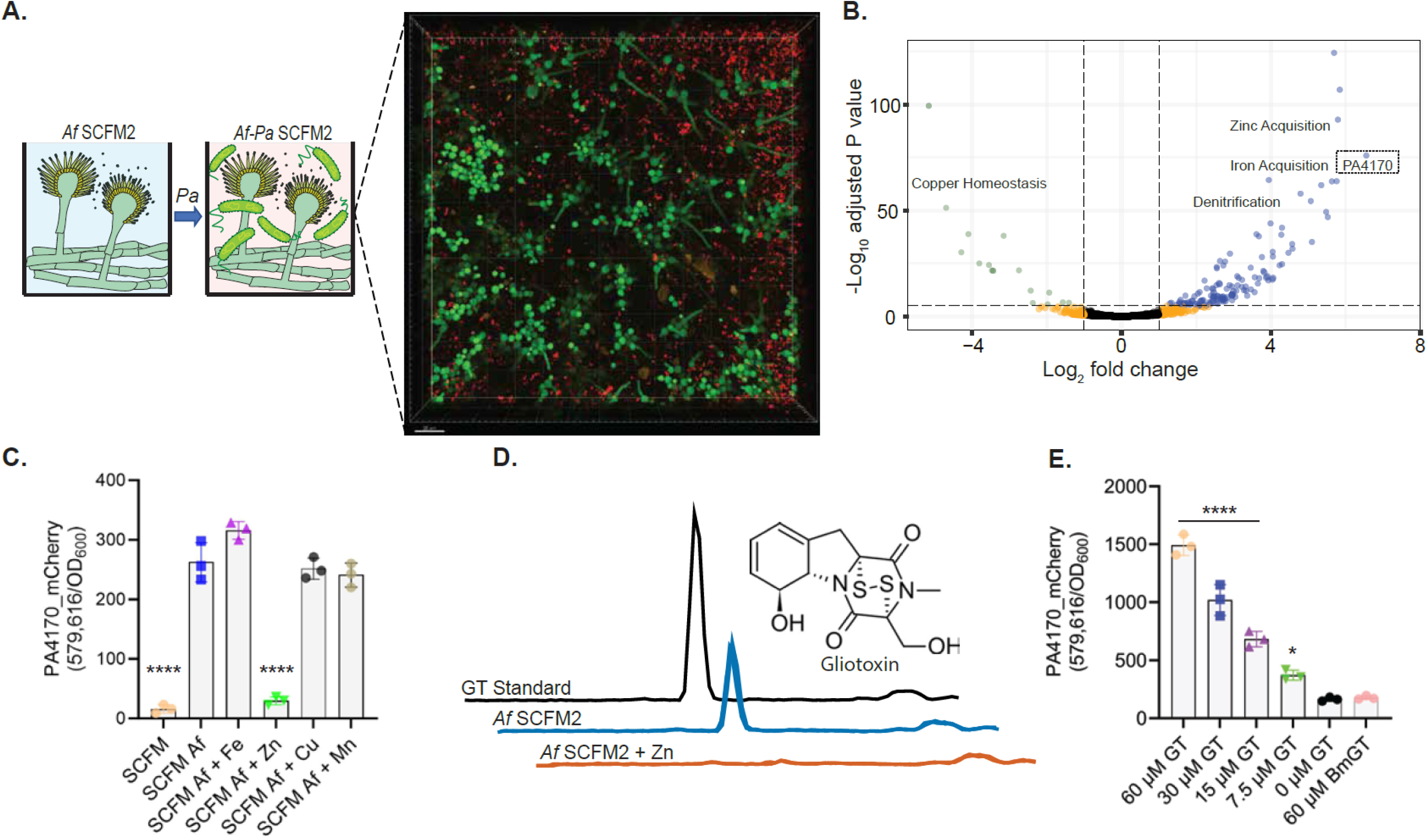
PA4170 is the most highly differentially expressed *P. aeruginosa* gene during co-culture with *A. fumigatus* in SCFM2, and expression is activated upon gliotoxin exposure. **A.** Schematic illustrating the *A. fumigatus* and *P. aeruginosa* co-culture setup. *A. fumigatus* is first inoculated into SCFM2 to facilitate germination and hyphal development, followed by *P. aeruginosa* inoculation. Co-culture was imaged by confocal laser scanning microscopy (CLSM) at 18 hr, 37 °C. *A. fumigatus* was labelled with constitutively expressed mNeonGreen, and *P. aeruginosa* with mCherry. The data are representative of three independent experiments, each performed in triplicate. Scale bar = 20 µm **B.** Volcano plot illustrating *P. aeruginosa* genes differentially expressed upon co-culture with *A. fumigatus*. Bubble colors are used to indicate genes significantly altered in expression with a Log_2_ fold change greater than 1 (blue), less than −1 (green), above the adjusted P-value threshold (orange) or unchanged (black). Significantly altered clusters of genes (>5) with established functions in *P. aeruginosa* are shown (zinc acquisition, iron acquisition, denitrification, copper homeostasis). PA4170 (dotted box) was the highest differentially expressed gene (93.7-fold-change). The experiment was performed using biological triplicates. **C.** mCherry-tagged PA4170 reporter assay examining PA4170 expression following *P. aeruginosa* growth in spent *A. fumigatus* SCFM culture supernatants (24 hr) supplemented with different metals (5 µM FeSO_4_, ZnSO_4_, CuSO_4_, MnSO_4_). Fluorescence was read at 6 hr and normalized to cell density (OD600). The data are representative of three independent experiments performed in triplicate. Statistical significance was calculated by one-way ANOVA with Dunnett’s multiple comparisons test using ‘SCFM *Af*’ as the control column (**** P < 0.0001). Error bars represent standard deviation from the mean. **D.** Untargeted metabolomics of *A. fumigatus* culture supernatants during growth in SCFM and SCFM + 10 µM ZnSO_4_ (24 hr) identified gliotoxin as significantly lowered upon zinc supplementation. Chromatograms show the peak corresponding to gliotoxin (triplicate), alongside a commercial standard. The experiment was performed using biological triplicates. **E.** mCherry-tagged PA4170 reporter fluorescence assay examining PA4170 protein expression in SCFM following gliotoxin (GT) or bisthiomethylgliotoxin (bmGT) supplementation (0 – 60 µM). Fluorescence was read at 8 hr and normalized to cell density (OD600). The data are representative of two independent experiments performed in triplicate. Statistical significance was calculated by one-way ANOVA with Dunnett’s multiple comparisons test using ‘0 µM GT’ as the control column (* P = 0.0197, **** P < 0.0001). Error bars represent standard deviation from the mean.

Upon co-culture with *A. fumigatus*, 217 *P. aeruginosa* genes increased and 53 decreased in expression compared to monoculture (fold change ≥ 2 and Padj < 0.05, **Dataset S1**), indicating that there is a robust *P. aeruginosa* response to the presence of *A. fumigatus*. **Fig. 1B** shows an overview of the differentially expressed genes. During growth in oxygen restricted environments, *P. aeruginosa* can anaerobically respire through the process of denitrification, using nitrate as a terminal electron acceptor. Although *A. fumigatus* can grow at low oxygen concentrations, this fungus cannot grow under strict anaerobic conditions and requires a functional respiratory chain for the initiation of infections (33). Many genes were part of well-characterized regulons involved in adaptation to oxygen levels and micronutrient (iron, copper, zinc) limitation **(Fig. 1B** and **Dataset S1**). Genes controlled by the Fnr-type transcription factors Anr and Dnr, which regulate the low-oxygen and denitrification networks in *P. aeruginosa* (34), were upregulated upon co-culture (**Dataset S1**). This suggests that the competition with *A. fumigatus* for oxygen drives *P. aeruginosa* to utilize alternative electron acceptors. Genes required for synthesis of the iron-scavenging siderophores pyoverdine and pyochelin were also upregulated (**Dataset S1**). *P. aeruginosa* central metabolic rewiring in response to iron limitation was also evident upon co-culture, with an increased expression of fumarate hydratase (*fumC1*, 27-fold higher) and superoxide dismutase (*sodM*, 15-fold higher) (**Dataset S1**). Genes required for zinc acquisition were also highly upregulated during co-culture (**Fig. 1B**, **Dataset S1**). This included genes required for synthesis of the zincophore pseudopaline, dedicated zinc uptake systems, and genes encoding zinc-independent ribosomal protein paralogs, which are known to be activated in *P. aeruginosa* upon zinc starvation (35, 36). *dksA2*, encoding a zinc-independent structural analog of the transcription factor DksA (37), was expressed 42-fold higher during co-culture, alongside zinc-independent ribosomal subunits. Genes involved in copper homeostasis were highly downregulated in expression during co-culture (**Fig. 1B**, **Dataset S1**), including c*opA1,* which encodes an ATPase expressed in response to high Cu (38), and the copper-regulated toxin fluopsin C (39). These results indicate that, during co-culture with *A. fumigatus* in a validated CF pre-clinical model, *P. aeruginosa* induces genes encoding proteins involved in low oxygen growth and acquisition of zinc and iron.

### PA4170 is activated by a zinc-regulated *A. fumigatus* metabolite

Although most of the differentially expressed genes upon co-culture with *A. fumigatus* could be assigned to known regulons, the top *P. aeruginosa* gene induced (93-fold) upon *A. fumigatus* exposure (**Fig. 1B**), PA4170 (gene designation PA14_09950 in *P. aeruginosa* strain PA14), was uncharacterized. PA4170 is a hypothetical protein, predicted to be encoded on a monocistronic mRNA, and previously shown to be transcriptionally induced during *P. aeruginosa* human chronic infection compared to in vitro mono-culture growth (26).

To explore the function of PA4170, we first set out to identify the *A. fumigatus* signal/cue that induces PA4170. To accomplish this, we generated a PA4170-mCherry fusion protein in *P. aeruginosa*, in which mCherry was fused to the C-terminus of the full-length protein. This facilitated the quantification of PA4170 production under different growth regimes using a high-throughput fluorescence assay. In agreement with our previous transcriptome results, PA4170 levels were low when *P. aeruginosa* was cultured alone in SCFM2 or standard laboratory media (**Fig. 1C, Fig. S1C**). As expected, PA4170 levels increased upon *P. aeruginosa* co-culture with *A. fumigatus* in SCFM2 (**Fig. S1C**), and cell-free spent culture supernatants from *A. fumigatus* grown alone also activated PA4170 production (**Fig. 1C**). The PA4170 inducer could be organically extracted from spent *A. fumigatus* culture supernatants with ethyl-acetate, suggesting that it was a non-polar small molecule (**Fig. S1C**). In addition, spent *A. fumigatus* culture supernatants from as early as 9-12 hr growth in SCFM2 were capable of inducing *P. aeruginosa* PA4170 expression (**Fig. S1D**).

Since SCFM2 is a low metal environment and *P. aeruginosa* was metal-limited during co-culture (**Fig. 1B**), we hypothesized that the PA4170 inducer was synthesized by *A. fumigatus* in response to metal limitation. To test this hypothesis, *A. fumigatus* was grown in SCFM2 in the presence of various metals (iron, zinc, manganese, copper), and supernatants from these cultures tested for the ability to induce PA4170-mCherry. While supplementation with iron, manganese, and copper had no impact on the ability of *A. fumigatus* supernatants to induce PA4170-mCherry, addition of zinc eliminated induction (**Fig. 1C**). Importantly, the addition of zinc and other metals to *A. fumigatus* culture supernatants obtained after growth in SCFM2 not supplemented with metals retained the ability to activate PA4170 (**Fig. S1E**), indicating that zinc addition does not impact the activity of the PA4170 inducer after synthesis. These data support zinc limitation as a key cue required for *A. fumigatus* synthesis of the PA4170 inducer.

### *A. fumigatus* gliotoxin induces PA4170

To identify *A. fumigatus* metabolites produced during zinc limitation, we performed untargeted metabolomics of *A. fumigatus* culture supernatants following growth in SCFM2 and SCFM2 + zinc. These data revealed that the *A. fumigatus* epipolythiodioxopiperazine (ETP) natural product gliotoxin was produced at high levels in SCFM2 but was undetectable in SCFM2 supplemented with zinc (**Fig. 1D, Fig. S1F**). ETPs are a class of disulfide-containing, toxic secondary metabolites made exclusively by fungi (40). Gliotoxin is the most well-studied member of the ETP class and is produced in a pro-drug form with an intact disulfide bond (**Fig. 1D**). The disulfide bridge is critical for the intracellular deleterious effects of gliotoxin, mediated through redox cycling and protein conjugation within target organisms (reviewed in (41)). Gliotoxin synthesis is controlled by multiple cues in *A. fumigatus*, including zinc starvation (42, 43), and global fungal secondary metabolism regulators such as LaeA (44). Therefore, it is likely that *A. fumigatus* is zinc-limited during growth in SCFM2, leading to the activation of gliotoxin biosynthesis. In support of this, production of the *A. fumigatus* zinc starvation regulated proteins Aspf2 (45) and GliT (42) were highly expressed during growth in SCFM2 (**Fig. S1G**), and zinc supplementation reduced their synthesis (**Fig. S1G**).

To test if gliotoxin induces PA4170 expression, *P. aeruginosa* carrying PA4170-mCherry was exposed to gliotoxin and mCherry levels quantified. PA4170 was activated by gliotoxin in a dose-dependent manner (**Fig. 1E**). As a control, *P. aeruginosa* carrying PA4170-mCherry was exposed to Bisdethiobis(methylthio)gliotoxin (BmGT), a S-thiomethylated, redox-inactive derivative of gliotoxin. BmGT did not activate expression of PA4170 (**Fig. 1E**) indicating that the gliotoxin disulfide bridge gliotoxin was crucial to activate PA4170 expression.

Spent culture supernatants of other *A. fumigatus* strains from clinical (AF293, AF210) and environmental sources (A1241) were then tested with the *P. aeruginosa* DnoP-mCherry reporter assay. All strains substantially induced DnoP expression (**Fig. S1H**). Spent culture supernatants from a Δ*gliZ* mutant (CEA10 background), which cannot synthesize gliotoxin (46), were greatly diminished in their ability induce DnoP expression (**Fig. S1H**). Zinc supplementation prior to *A. fumigatus* inoculation obliterates this response (**Fig. S1I**). These data indicate that gliotoxin is produced by diverse *A. fumigatus* isolates in SCFM2 in response to zinc limitation, and this metabolite induces expression of PA4170 in *P. aeruginosa*.

### Gliotoxin induces PA4170 production by zinc starvation

ETPs are thought to enter cells where it is converted a reduced, active form which can chelate zinc with high affinity (47, 48), thus inhibiting growth. To explore the response of *P. aeruginosa* to gliotoxin in more detail, we carried out an RNA-seq experiment of *P. aeruginosa* exposed to gliotoxin (30 µM) during growth in SCFM2 compared to a solvent control. Gliotoxin exposure resulted in 219 *P. aeruginosa* genes increased and 81 decreased in expression (fold change ≥ 2 and Padj < 0.05, **Dataset S1**). 29% of the genes differentially expressed in the *P. aeruginosa*-*A. fumigatus* co-culture experiment (**Fig. 1A**) were also differentially regulated by gliotoxin, indicating that gliotoxin is an important contributor to the *P. aeruginosa* transcriptome during co-culture. Examination of the differentially regulated genes revealed that gliotoxin exposure elicits a specific, zinc starvation response in *P. aeruginosa*, with PA4170 as the most upregulated transcript (313-fold) (**Fig. 2A**). Several of these upregulated genes (50) are part of the Zur regulon, which are induced by zinc starvation (**Dataset S1**), suggesting that PA4170 induction is mediated by zinc starvation. To further test this, *P. aeruginosa* containing the PA4170-mCherry fusion protein was exposed to N,N,N’,N’-tetrakis-(2-Pyridylmethyl) ethylenediamine (TPEN). TPEN is a membrane-permeable, heavy metal chelator with high affinity for zinc. Exposure of *P. aeruginosa* to TPEN activated PA4170 production in a dose-dependent manner, and this activation could be suppressed by zinc supplementation (**Fig. S2A**). These data indicate that ETP-mediated zinc chelation specifically activates PA4170 as part of the *P. aeruginosa* response to zinc depletion, a response downstream of, or parallel to, the Zur regulon.

**Fig. 2.**
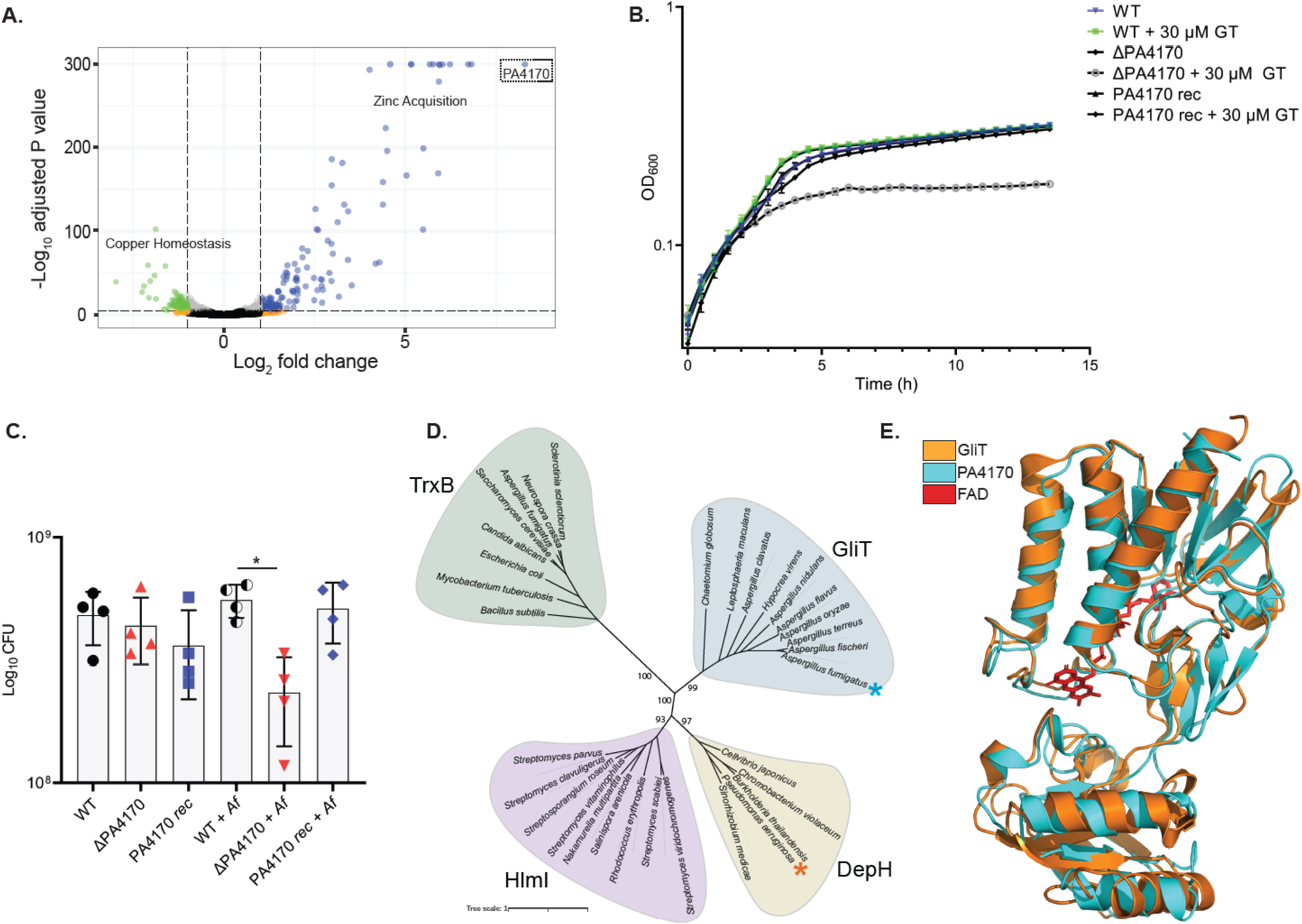
PA4170 encodes a dithiol oxidase which protects *P. aeruginosa* against disulfide-containing natural products. **A.** Volcano plot illustrating *P. aeruginosa* genes differentially expressed upon exposure to gliotoxin (20 µM) versus a solvent (MeOH) control. Bubble colors are used to indicate genes significantly altered in expression with a Log_2_ fold change greater than 1 (blue), less than −1 (green), above the adjusted P-value threshold (orange) or unchanged (black/grey). Significantly altered groups of genes (>5) with established functions in *P. aeruginosa* are shown (zinc acquisition, copper homeostasis). PA4170 (dotted box) was the highest differentially expressed gene (313-fold-change). The experiment was performed using biological triplicates. **B.** Growth of a ΔPA4170 deletion mutant is more susceptible to gliotoxin exposure (30 µM) when compared to the *P. aeruginosa* wild type or PA4170 complemented strain (PA4170 rec). Strains cultured in M9 succinate. An equivalent volume of solvent (DMSO) was added to control samples. The data are representative of three independent experiments performed in triplicate. **C.** Growth of a ΔPA4170 deletion mutant is reduced upon *A. fumigatus* co-culture when compared to wild type *P. aeruginosa* during co-culture. Growth of ΔPA4170 is not significantly altered compared to the wild type or PA4170 rec (PA4170 plasmid complemented) in monoculture. Cultures were plated for CFU counts 16 hr after *P. aeruginosa* addition. CFU = colony-forming unit. The data are representative of three independent experiments performed in quadruplicate. Statistical significance was calculated by Brown-Forsythe ANOVA test with Dunnett’s T3 multiple comparisons (*P = 0.0215). Error bars represent standard deviation from the mean. **D.** Phylogenetic analysis of PA4170 alongside representative thioredoxin oxidoreductases (TrxB) and dithiol oxidases from various fungi (GliT) and bacteria (DepH, Hlml). Amino acid sequences were obtained from GenBank database and aligned with ClustalW. Representative bootstrap values are shown. **E.** Overlay (PyMOL) of GliT X-ray structure (PDB: 4NTC, Orange) with the high-confidence PA4170 AlphaFold (71) predicted structure (Cyan) indicates remarkable structural similarity between these two proteins. Flavin adenine dinucleotide (FAD), which was co-crystallized bound in the GliT active site is shown in red.

### PA4170 plays a role in *P. aeruginosa* ETP and Dithiolopyrrolone (DTP) resistance

To functionally characterize PA4170, a *P. aeruginosa* ΔPA4170 deletion mutant was generated. ΔPA4170 had no discernible growth phenotype during growth in SCFM2 or other laboratory media (**Fig. 2B**). However, this strain was significantly more susceptible to gliotoxin exposure (30 µM) when compared to the wild type (**Fig. 2B**), and this hyper-susceptibility could be genetically complemented by introduction of PA4170 into ΔPA4170. ΔPA4170 was not hyper-susceptible to bmGT exposure (**Fig. S2B**), or the zinc chelator TPEN (**Fig. S2C**), ruling out a direct role for this protein in *P. aeruginosa* zinc homeostasis. ΔPA4170 also exhibited a growth defect when cultured alongside *A. fumigatus* in SCFM2, compared to wild type *P. aeruginosa* (**Fig. 2C**). This growth defect was not evident when ΔPA4170 was cultured alongside a Δ*gliZ* mutant (**Fig. S2D**). These data indicate that PA4170 is critical for *P. aeruginosa* protection to gliotoxin exposure and fitness during co-culture with *A. fumigatus*.

Does PA4170 provide protection against ETPs other than gliotoxin? To answer this question, we exposed wild-type *P. aeruginosa* and ΔPA4170 to four additional dithiol-containing microbial-produced toxins, including three ETP toxins (chaetocin, chetomin, and romidepsin) and one Dithiolopyrrolone (DTP) toxin (holomycin). DTP toxins are produced by bacteria and similar to ETPs, impact metal homeostasis in both prokaryotic and eukaryotic microbes (47, 49). Growth of ΔPA4170 was reduced upon exposure to chaetocin, chetomin, and holomycin, but not romidepsin, compared to WT *P. aeruginosa* (**Fig. S3A-D**). In addition, exposure to chaetocin, chetomin and holomycin activated *P. aeruginosa* PA4170 expression (**Fig. S3E**). These data indicate that PA4170 provides protection against a range of microbial produced dithiol-containing toxins.

### PA4170 shares catalytic motifs with bacterial and fungal natural product dithiol oxidases

The role of PA4170 in protection against dithiol toxins motivated us to examine the phylogeny of PA4170 in more detail. PA4170 was found in most fully sequenced *P. aeruginosa* strains (279/284), as well as a small number of other pseudomonads including *P. fluorescens*, *P. chlororaphis*, *P. putida*, *P. stutzeri,* and *P. azotoformas* isolates. Although PA4170 homologs appear to be distributed at random across non-*P. aeruginosa* pseudomonads, it is notably present in all sequenced isolates within a particular *P. chlororaphis* clade (**Fig. S4**).

An InterPro functional prediction indicated that PA4170 encodes a flavin adenine dinucleotide (FAD) flavoprotein belonging to the family of pyridine nucleotide-disulfide oxidoreductases. These enzymes contain a characteristic pair of redox-active cysteines involved in the transfer of reducing equivalents from the FAD cofactor to a substrate. NCBI PDB-BLAST was used to scan the Protein Data Bank repository for structurally elucidated, distantly related proteins with sequence similarity to PA4170. Three of the nine proteins identified in this analysis are functionally characterized and were shown to catalyze disulfide bond formation in natural products (**Fig. S5A-C**) including: DepH from *Chromobacterium violaceum* (50), HlmI from *Streptomyces clavuligerus* (51), and GliT from *A. fumigatus* (52), which catalyze disulfide bridge closure of romidepsin, holomycin, and gliotoxin respectively.

A phylogenetic analysis of PA4170 and other thioredoxin oxidoreductase-like proteins from various fungi and bacteria revealed a thioredoxin oxidoreductase (TrxB) group, and three separate groups where the structurally elucidated HlmI, GliT, and DepH are representative members (**Fig. 2D**). All enzymes share the conserved CXXC motif which is essential for dithiol oxidase activity (50, 51, 53, 54). The GliT group consists of fungal proteins required for the biosynthesis of ETPs, and it likely diverges from the HlmI and DepH groups because of its fungal origin. PA4170 falls within the DepH group, which also contains TdpH from *Burkholderia thailandensis* E264, known to catalyze disulfide bond formation in thailandepsin A and B biosynthesis (55). While the BLAST analysis indicated a low sequence identity (28.6%) between PA4170 and GliT (**Fig. S5B**), a structural overlay of an AlphaFold generated structural model of PA4170 with the known crystal structure of GliT demonstrated remarkable structural similarity, with conservation at many key motifs and residues known to be required for ETP and DTP oxidation (**Fig. 2E**, **Fig. S6**). These data suggest that PA4170 protected *P. aeruginosa* against ETP and DTP toxin exposure through disulfide bond regeneration, precisely the same mechanism used by *A. fumigatus* for GliT-mediated self-protection from gliotoxin.

### PA4170 is a <u>D</u>isulfide-containing <u>N</u>atural product <u>O</u>xidase in <u>P</u>seudomonas (DnoP) with broad substrate specificity

Intrigued by the ability of this single enzyme to protect *P. aerugino*sa against multiple disulfide-containing natural products (**Fig. S3A-D**), we investigated the enzymology of PA4170 in more detail. Both GliT and PA4170 were recombinantly expressed in *E. coli*, purified to homogeneity, and assessed for their ability to oxidize the ETP/DTP toxins gliotoxin, chaetocin, chetomin, romidepsin, and holomycin. Recombinant PA4170 and GliT catalyzed disulfide oxidation of both gliotoxin and holomycin (**Fig. 3AB**). However, PA4170 was also capable of restoring the disulfide bridge of the reduced ETPs chetomin and chaetocin, whereas GliT was unable to act on these metabolites. Neither enzyme was capable of oxidizing reduced romidepsin (**Fig. 3A-C**). Based on these data, we herein assigned *P. aeruginosa* PA4170 as <u>D</u>isulfide-containing <u>N</u>atural Product <u>O</u>xidase, *<u>P</u>seudomonas* (DnoP). From examining the bulk *P. aeruginosa* transcriptome from CF patients colonized with this microbe across different CF clinics, we uncovered that the *dnoP* transcript is expressed highly during human infection (19, 20, 23). However, *P. aeruginosa* chronic infection isolates are highly diverse, so although expression of the *dnoP* transcript was detected, these individual isolates may not make a functional DnoP protein. To address this question, we modified our dithiol oxidase enzyme assay to test the DnoP activity within *P. aeruginosa* cell lysates. Diverse *P. aeruginosa* isolates (PA14, PA14 Δ*dnoP*, PAO1, C5912M and C2773C) were treated with gliotoxin (or a solvent control) during growth in SCFM, and the resulting cell lysates were tested for DnoP activity. As shown in Fig. 3D, inducible DnoP activity was readily detected in diverse *P. aeruginosa* isolates. C5912M is a mucoid *P. aeruginosa* isolate from a patient with CF (56). This result suggests that mucoidy, which is known to impact antimicrobial efficacy (33), does not impact the ability of gliotoxin to activate *dnoP* expression. Importantly, gliotoxin exposed cell lysates from the PA14 Δ*dnoP* mutant had only background levels DnoP enzyme activity. These data conclusively show that DnoP functionality is conserved across *P. aeruginosa* isolates.

**Figure 3.**
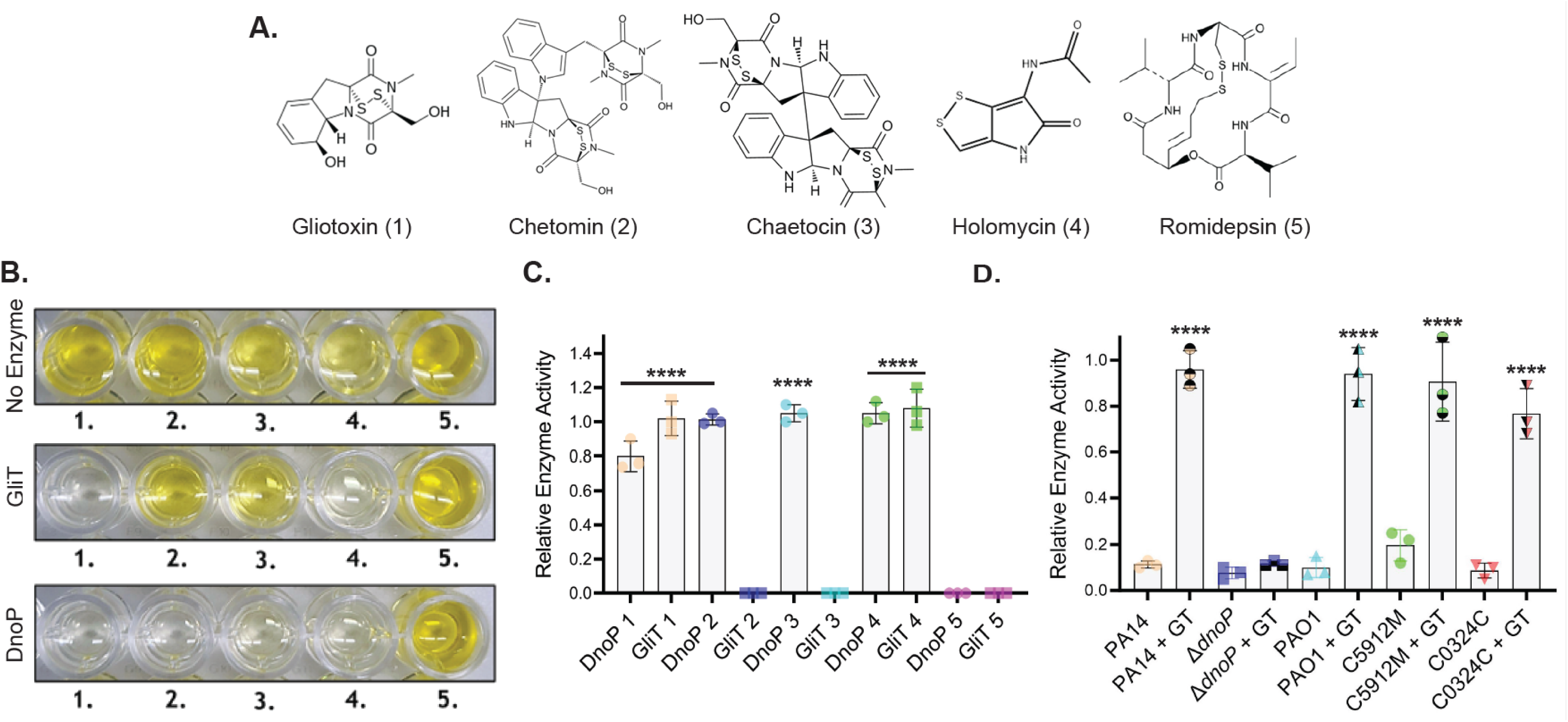
DnoP can catalyze disulfide bond formation for diverse disulfide-containing natural products. **A.** Structures of the disulfide-containing natural products 1; gliotoxin (*Aspergillus fumigatus*), 2; chetomin (*Chaetomium cochliodes*), 3; chaetocin (*Chaetomium minutum*), 4; holomycin (*Streptomyces clavuligerus*) and 5; romidepsin (*Chromobacterium violaceum*). **B.** DTNB (5,5’-dithio-bis-(2-nitrobenzoic acid)) assay image demonstrating that recombinantly expressed GliT and DnoP can oxidize both reduced gliotoxin (1) and reduced holomycin (4), resulting in an absence of yellow-colored 2-nitro-5-thiobenzoate (TNB) formation upon DTNB addition, due to the enzyme mediated oxidation of the natural product free sulfhydryl groups. DnoP is also capable of oxidizing reduced chetomin (2) and reduced chaetocin (3), whereas neither DnoP nor GliT can oxidize reduced romidepsin (5). Relative enzyme activity (the reduction in TNB formation at 412 nm relative to no-enzyme controls) for DnoP and GliT with the TCEP reduced substrates 1-5 is shown in **C**. 0.2 µM of DnoP/GliT enzyme was used in each assay. Reactions were incubated for 15 min at RT, followed by DTNB addition for 5 min before reading. The data are representative of three independent experiments performed in triplicate. The third replicate was performed with 0.2 µM GliT-mCherry and DnoP-mCherry purified from *A. fumigatus* (Fig. S7B). Statistical significance was calculated by one-way ANOVA with Dunnett’s multiple comparisons test using no enzyme added as the control (**** P < 0.0001). Error bars represent standard deviation from the mean. **D.** Relative DnoP enzyme activity (the reduction in TNB formation at 412 nm relative to no-enzyme controls) of *P. aeruginosa* whole cell lysates from PA14, Δ*dnoP*, PAO1, C5912M and C0324C, using TCEP reduced gliotoxin as a substrate. DnoP enzyme expression was induced through the addition of gliotoxin (20 µM) and compared to the addition of a solvent control (MeOH). Reactions were incubated for 15 min at RT, followed by DTNB addition for 5 min before reading. The data are representative of three independent experiments performed in triplicate. Statistical significance was calculated by one-way ANOVA with Dunnett’s multiple comparisons test using ‘PA14’ as the control column (**** P < 0.0001). Error bars represent standard deviation from the mean.

### DnoP functionally complements *A. fumigatus* Δ*gliT*

At this point, it was unclear whether the broader substrate specificity of DnoP results in a significant cost to catalytic function. To explore this, we tested the ability of heterologously expressed DnoP to substitute for the combined biosynthetic and self-defense functionalities of GliT in *A. fumigatus*. An *A. fumigatus* Δ*gliT* mutant was generated, which does not produce gliotoxin due to an inability to catalyze the final disulfide bridge closure. This mutant is also susceptible to exogenous gliotoxin exposure (52). Using CRISPR-Cas9, *A. fumigatus* Δ*gliT* was genetically complemented by placement of a DNA fragment consisting of the *gliT* native promoter and the *gliT* ORF with an in-frame C-terminal mCherry tag (*gliT*_mC) into an intergenic “safe haven” region of the *A. fumigatus* genome. The same procedure was used create another strain in which a DNA fragment containing the *P. aeruginosa* DnoP ORF with a C-terminal mCherry tag (*dnoP*_mC), driven by the *gliT* promoter, was inserted into *A. fumigatus* Δ*gliT*. As expected, complementation of Δ*gliT* with *gliT*_mC restored self-protection against gliotoxin to wild-type levels (**Fig. 4A**). Complementation of Δ*gliT* with *dnoP*_mC also restored gliotoxin self-protection, in a manner indistinguishable to the native enzyme (**Fig. 4A**). Furthermore, both *gliT_mc* and *dnoP_mc* complemented Δ*gliT* had gliotoxin biosynthesis fully restored upon growth in SCFM2 (**Fig. 4B**) and spent culture supernatants from these complemented strains activated DnoP expression in *P. aeruginosa* (**Fig. 4C**). Finally, purified GliT-mC and DnoP-mC (**Fig. S7B**) exhibited identical substrate specificity to the His-tagged versions of these proteins recombinantly expressed in *E. coli* (**Fig. 3C**). These data indicate that DnoP can effectively substitute for *A. fumigatus* GliT functionality, despite its low protein sequence identity and broad substrate specificity.

**Fig. 4.**
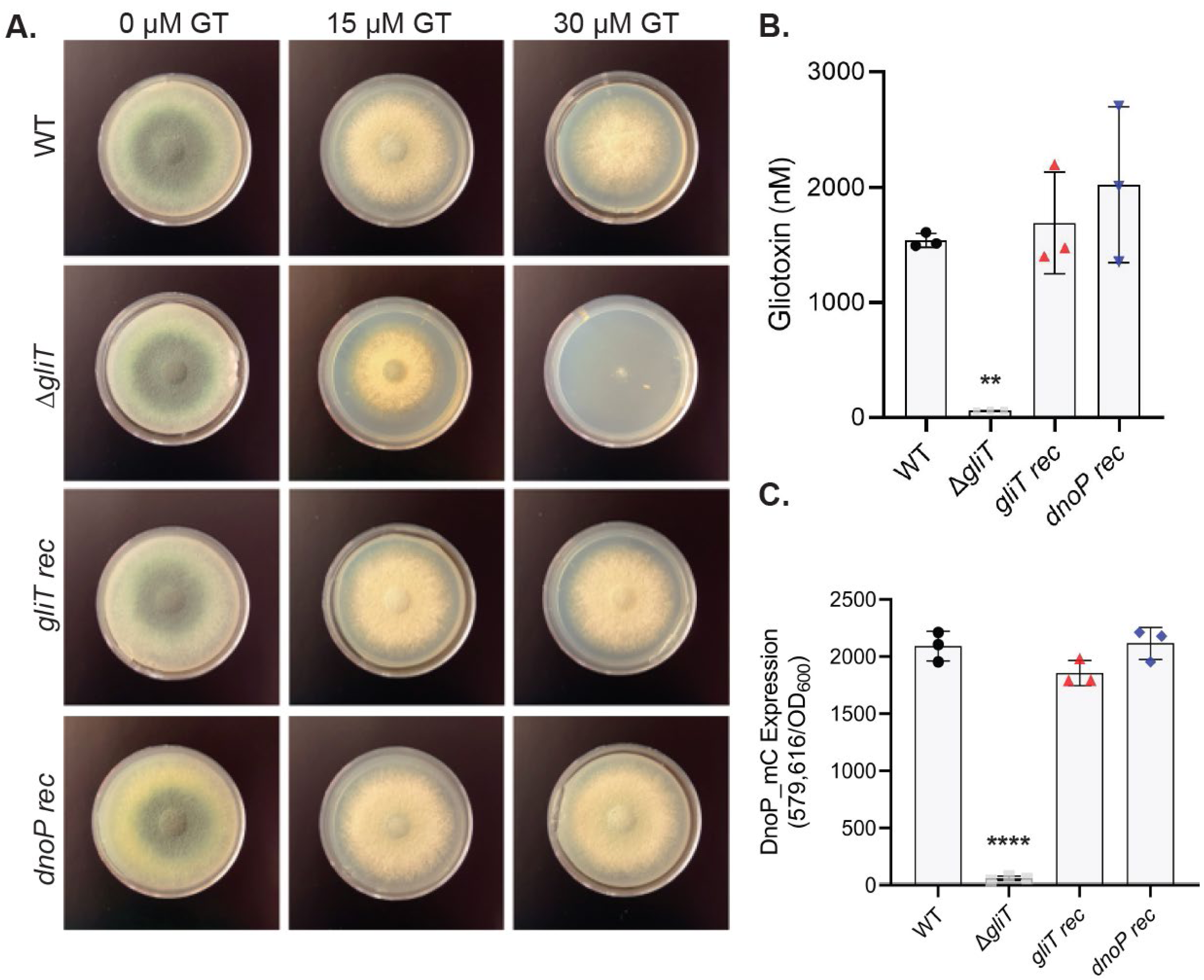
Heterologous expression of *dnoP* in *A. fumigatus* can functionally complement for the loss of the dithiol gliotoxin oxidase *gliT*. A. Plate assay of *A. fumigatus* wild type, Δ*gliT*, *gliT rec* and *dnoP rec* growth upon gliotoxin addition (0, 15, 30 µM) to GMM agar plates. Plates imaged at 48 hr. The data are representative of three independent experiments performed in triplicate. **B.** Complementation of Δ*gliT* in a neutral ‘safe-haven’ site with *gliT* (*gliT rec*) or *dnoP* (*dnoP rec*) restores *A. fumigatus* gliotoxin production to wild type levels. Gliotoxin was quantified using LC-MS following 72 hr growth in SCFM2. The experiment was performed in triplicate. Statistical significance was calculated by one-way ANOVA with Dunnett’s multiple comparisons test using ‘WT’ as the control column (** P = 0.0052). Error bars represent standard deviation from the mean. **C.** Complementation of Δ*gliT* in a neutral site with *gliT* (*gliT rec*) or *dnoP* (*dnoP rec*) restores the ability of *A. fumigatus* spent culture supernatants (24 hr) to activate *P. aeruginosa* PA4170 expression. Fluorescence was read at 10 hr and normalized to cell density (OD600). The data are representative of two independent experiments performed in triplicate. Statistical significance was calculated by one-way ANOVA with Dunnett’s multiple comparisons test using ‘WT’ as the control column (**** P < 0.0001). Error bars represent standard deviation from the mean.

## Discussion

Many microbes exist in natural environments as complex, interkingdom polymicrobial communities. Consequently, microbes have evolved diverse mechanisms to compete with their neighbors for space and resources. Nutritional resources are at the forefront of microbial competition, and microbes have evolved numerous strategies to increase nutrient acquisition or to actively restrict them from others (57, 58). We have identified and characterized a *P. aeruginosa* disulfide-containing natural product oxidase (DnoP) engaged in xenobiotic defense against a family of potent small molecule toxins that function by zinc sequestration. Importantly, DnoP was uncovered in *P. aeruginosa*, an organism which does not synthesize disulfide-containing natural products. All other dithiol oxidases characterized to date are encoded within natural product biosynthetic gene clusters, dedicated primarily to the synthesis of these toxins. Although the natural substrate of DnoP is uncertain, we uncovered that *dnoP* expression is activated upon co-culture with *A. fumigatus*, and that a Δ*dnoP* mutant has a fitness defect during co-culture with this ubiquitous, saprophytic fungus. Overall, the widespread conservation of *dnoP* in diverse *P. aeruginosa* isolates suggests that this bacterium frequently encounters ETP-producing filamentous fungi or DTP-producing bacteria in nature and has evolved a single protein system to defend against their toxins. It is likely that the broader role of ETPs and other dithiol-containing natural products in microbial interactions has been masked by the presence of cognate resistance genes like *dnoP*.

Microbial stress responses which detect ecological competition, such as nutrient limitation, are often coupled with the release of toxins, as this form of competition normally implies the presence of foreign genotypes. Nevertheless, it is unclear why the limitation of one nutrient but not another should promote toxin production (59). We propose that, as ETPs dysregulate zinc homeostasis in target organisms, orchestrating gliotoxin production via zinc starvation-responsive transcriptional regulators allows *A. fumigatus* to synthesize this toxin when it will have maximum impact on target organisms. The efficacy of an intracellular zincophore is likely to have maximum impact on target cells when the capacity of these competitors to obtain zinc is limited. Alternatively, it may be preferential for *A. fumigatus* to synthesize gliotoxin when its metabolism is transcriptionally wired to endure zinc-limitation, as microbes are acutely sensitive to shifts in cellular free zinc levels. It is likely that ETP biosynthesis requires a careful coordination between primary and secondary metabolism to avoid self-intoxication. Orchestrating ETP biosynthesis through a zinc starvation response may also facilitate cooperation in neighboring ETP producing fungi, catalyzing a positive feedback loop of toxin production (60). This fundamental regulatory cue may be conserved across multiple dithiol toxin producing organisms. Indeed, zinc starvation was shown to activate holomycin production in the Gram-negative marine bacterium *Photobacterium galatheae* (61).

Stress responses are an effective cue to provide a finely tuned, graded readout of ecological competition and its capacity to reduce cellular fitness. One which is receptive to both alterations in community species composition and to variations in the chemical signatures of competitors (59). In an example of cross-kingdom cue convergence, we have shown that *P. aeruginosa* expresses DnoP in response to zinc-starvation. The precise signaling cascade leading to *dnoP* activation has yet to be elucidated. However, this method of transcriptional regulation may provide *P. aeruginosa* pre-emptive protection against ETP exposure during growth in zinc-limited environments, where gliotoxin production by *A. fumigatus* is maximal. Alternatively, the activation of *dnoP* by zinc-starvation may allow for sustained *dnoP* expression upon ETP mediated disruption of cellular zinc homeostasis, facilitating survival. Thus, zinc limitation may be a conserved signal across kingdoms for both dithiol-containing natural product production and defense.

The ability of *dnoP* to fully complement for *gliT* functionality in *A. fumigatus* was unexpected. This suggests that there is minimal catalytic tradeoff in retaining a broader substrate range with these enzymes. Specialization is driven by requirements for adequate regulation and localization, as well as for high catalytic efficiency with the target substrate (62). The substrate specialization commonly found in dedicated natural product dithiol oxidases implies that the broad substrate specificity of DnoP is by design. It will be interesting to further explore the substrate specificity of DnoP from other organisms to see if this is a pervasive trend. In addition to providing evolutionary insight, these promiscuous enzymes may yield useful starting material for protein engineering and synthetic biology approaches.

In sum, this work provides strong evolutionary evidence that *P. aeruginosa* and other *dnoP*-containing pseudomonads are exposed to the antimicrobial activities of dithiol-containing toxins in natural settings. While this work leveraged a benchmarked clinical environment, it’s likely the conserved and tight coupling of dithiol-containing toxin expression and defense to zinc availability drives the specificity of this fungal-bacterial interaction in a variety of terrestrial ecosystems.

## Acknowledgements

The metabolomic studies were supported by Georgia Institute of Technology’s Parker H. Petit Institute for Bioengineering and Bioscience including the Systems Mass Spectrometry Core Facility. We thank Sean Capps for assistance with confocal microscopy. This study was supported by grants from the Cystic Fibrosis Foundation (WHITEL20A0 to MW) and the Shurl and Kay Curci Foundation (to MW). S.K.D. was supported by a Herchel Smith Postdoctoral Fellowship and a Cystic Fibrosis Postdoctoral Fellowship (DOLAN20F0).

## Materials and Methods

### Media and growth conditions

*P. aeruginosa* cells were routinely cultured in lysogeny broth (LB LENNOX) at 37 °C with shaking (250 rpm) unless otherwise specified. All liquid culture experiments were performed in M9-succinate or SCFM unless explicitly stated otherwise. *Pseudomonas* isolation agar (PIA) was used for selecting *P. aeruginosa* against *A. fumigatus* for CFU plating. *Escherichia coli* cells were cultured in lysogeny broth (LB). 1.5% (wt/vol) agar was used to prepare plates. Antibiotics were added to the media as necessary: 60 µg ml−1 of gentamicin (Gen), 75 µg ml−1s tetracycline (Tet), or 300 µg ml−1 carbenicillin (Cb) for *P. aeruginosa*; 10 µg ml−1 of gentamicin (Gen), 10 µg ml−1 tetracycline (Tet), or 100 µg ml−1 carbenicillin (Cb) for *E. coli*. Cell growth was monitored as optical density in a spectrophotometer at a wavelength of 600 nm (OD_600_).

*Aspergillus fumigatus* and other *Aspergillus* isolates were stored as conidia in 25% glycerol at −80°C and maintained on 1% glucose minimal medium [GMM]. Solid medium was prepared by addition of 1.5% agar. All experiments were performed with GMM, SDB (Sabouraud-dextrose broth) or SCFM unless explicitly stated otherwise. For all experiments, *A. fumigatus* was grown on solid GMM at 37°C for 3 days to produce conidia. Conidia were collected using 0.01% Tween 80, counted using a hemocytometer, and diluted in medium to the final concentration used in each assay.

### *A. fumigatus* and *P. aeruginosa* SCFM co-culture model

*A. fumigatus* was incubated statically at 37°C in 5 ml SCFM2 (10^5^ conidia/mL), in 6-well flat bottom cell culture plates until germination and hyphal development (18 hr). *P. aeruginosa* was precultured in LB overnight, then 50 μl of the culture was inoculated into 5 ml of SCFM and incubated the culture at 37°C for 6 h until the cultures reached mid-log phase (optical density at 600 nm [OD_600_] = 0.5). The OD_600_ was adjusted to 0.001 in 5 ml of SCFM2, and this 5 ml was added to the germinated *A. fumigatus* culture to establish the co-culture, or into 5 ml SCFM2 for *P. aeruginosa* monoculture (*P. aeruginosa* starting OD_600_ = 0.0005 (∼5 × 10^5^ cells mL^−1^)). *P. aeruginosa* CFU was tracked over time by plating on PIA. Cultures were grown until *P. aeruginosa* reached stationary phase (12-14 hr). *A. fumigatus* mycelial wet weight (mg/mL^-1^) was quantified by filtering cultures containing *A. fumigatus* through a 40µm cell strain and rising with PBS. Co-culture biofilm images were acquired with a Zeiss LSM 880 CLSM utilizing Zen image capture software. Detection of mCherry-expressing *P. aeruginosa* cells was performed with an excitation wavelength centered at 587 nm and an emission wavelength centered at 610 nm. Detection of mNeonGreen-expressing *A. fumigatus* was performed using an excitation wavelength centered at 488 nm and an emission wavelength centered at 509 nm. All images were acquired using a 63× oil-immersion objective.

### *P. aeruginosa* and *A. fumigatus* co-culture RNA-Seq

Whole cultures (biofilm and planktonic cells) of *P. aeruginosa* monocultures and co-cultures with *A. fumigatus* were harvested in the stationary phase of growth (triplicates) by aspirating the contents of the petri-dish well into a 15 ml falcon tube (passed over a 40 µm cell strainer) and combining this with an equal volume of RNA-later. Cultures stored in RNAlater were pelleted, resuspended in 1 mL RNA Bee, and RNA was extracted as described previously. Sequencing was performed by the Microbial Genome Sequencing Center (https://www.migscenter.com/). Default parameters were used for all software. Trimmed reads were then mapped to *P. aeruginosa* PA14 reference (available for download from pseudomonas.com) genome using Bowtie2 v2.4.2 with default parameters for end-to-end alignment. Read summarization was performed using featureCounts (63). DESeq2 was employed to analyze differentially-expressed genes (64). Annotations of differentially expressed genes were obtained from the reference annotation of the *Pseudomonas* genome available at the PGD website. Genes were considered as significantly altered when their adjusted-P value was <0.05 (Dataset S1). The sequencing data are deposited at ArrayExpress (accession number E-MTAB-12786).

### *P. aeruginosa* GT exposure RNA-seq

*P. aeruginosa* PA14 was precultured as described above, and the OD_600_ was adjusted to 0.05 in 8 ml of SCFM2, in 6-well flat bottom cell culture plates. Gliotoxin (20 µM) or MeOH (solvent control) was added at log phase (optical density at 600 nm [OD_600_] = 0.3), and the cultures were incubated statically for an additional 4 h. 5 ml of each culture (triplicate) was then immediately added to an equal volume of RNA-later. RNA isolation, sequencing and mapping was carried out as described above. The sequencing data are deposited at ArrayExpress (accession number E-MTAB-12785).

### Statistics

All experiments were performed independently at least three times (unless otherwise stated) with similar observations. The exact *n* values for experiments are provided in figures and legends. Statistical analyses were conducted using Prism GraphPad 10 and specified in figure legends. Unless otherwise noted, all experiments were performed with a minimum of three biologically independent samples.

### PA4710-mCherry reporter assay

The *P. aeruginosa* PA4170-mCherry fusion protein reporter strain was sub-cultured to an OD_600_ of 0.05 in SCFM and 20 µl was then added to a 96-well plate wells containing *A. fumigatus* spent culture supernatants or SCFM containing the indicated supplements (180 µl). The resulting plates were incubated statically at 37°C for the indicated times (6-12 hr). Optical density (OD600) and fluorescence was measured in a BioTek SynergyMx plate reader (excitation, 587 nm; emission, 620 nm) with Gen5 software.

### *A. fumigatus* mCherry reporter assays

*A. fumigatus* mCherry fusion protein strains were inoculated into SCFM (10^5^ conidia/mL – 10 ml) into 6-well flat bottom cell culture plates. Plates were incubated statically at 37°C for 48 hr. Florescence was measured in a BioTek SynergyMx plate reader (excitation, 587 nm; emission, 620 nm) with Gen5 software. GliT-mCherry and Aspf2-mCherry florescence was normalized to *gpdA*p-grown in identical culture conditions to account for any growth variations.

### *A. fumigatus* gliotoxin exposure plate assays

Radial growth assays were performed by inoculating GMM agar plates containing the indicated quantity of gliotoxin with 10^3^ conidia in 5 μL 0.01% Tween 20, followed by incubation at 37°C for 48 h and imaging.

### LC-MS analysis of gliotoxin

Quantification of gliotoxin production was carried out by LC-MS as described previously (65). Briefly, culture supernatants from *A. fumigatus* grown for the indicated duration (24-72 hr) in SCFM2 were harvested and stored at −20°C. 100 µl of samples was added into a new tube, then 200ul of ice cold MeOH: ACN:H_2_O (5:3:2/V:V:V) was added. Samples were vortexed twice, centrifuged at 21,100x g for 5 min, and then 100 µl of sample was added to an LC vial. Ultra-Performance Liquid Chromatography-mass spectrometry (UPLC-MS) was performed using an UltiMate 3000 fitted with a Waters AcquityUPLC BEH C18 column (100 x 2.1 mm, 1.7 micron) coupled to Orbitrap ID-X mass spectrometry system.

The chromatographic method for sample analysis involved elution with H_2_O and 0.1% formic acid (mobile phase A) acetonitrile (ACN) and 0.1% formic acid (mobile phase B) at 0.4 ml min^−1^ flow rate using the following gradient program: 0 min 5% B; 4 min 30% B; 7min 100% B; hold to 8.9 min, then 9 min 5% B hold to 10 min. The column temperature was set to 50 °C, and the injection volume was 5 µL. The targeted molecule gliotoxin fragmented by HCD at 35 collision energy in positive mode. MS/MS transitions for gliotoxin are 327.05/263.08. Standard curves of gliotoxin were generated. The amount of gliotoxin in the samples was calculated based on the standard curve.

### Phylogenetic analysis of PA4170

BLASTN and BLASTP searches were performed on pseudomonas.com and NCBI to identify orthologs of PA4170 across the *Pseudomonas* genus and other organisms. DNA sequences encompassing the PA4170 locus from the *P. aeruginosa* PA14 strain was used as the query. Orthologs identified from complete bacterial genomes with E values of <1 × 10^−4^, percent identity >90%, and query coverage values of >90% were retained for further analysis. Sequences of PA4170 orthologs were aligned using ClustalW. The alignments were visualized in Jalview (66) using the default nucleotide color scheme. For the phylogenetic analysis of PA4170 alongside representative thioredoxin oxidoreductases (TrxB) and dithiol oxidases from various fungi (GliT) and bacteria (DepH, Hlml). Amino acid sequences were obtained from GenBank database and aligned with ClustalW. Phylogenetic analysis was performed using maximum likelihood methods (MEGA11). Representative bootstrap values were calculated and the tree was visualized and annotated in iTOL (67). A maximum-likelihood phylogenetic tree for *Pseudomonas* was constructed in MEGA (68) using 358 16s rRNA sequences representing diverse isolates. For better visualization of the tree, we only included approximately 50 16s sequences from the *P. aeruginosa* strains. The 16s sequences were aligned with ClustalW and used as an input in MEGA11. The tree was visualized and annotated in iTOL (67).

### Plasmids and strains

Strains and plasmids used in this study are listed in Supplementary Table 1. All plasmids were verified by PCR, restriction enzyme digestion, and if necessary, DNA sequencing. Verified plasmids for targeting *P. aeruginosa* were transformed into *E. coli* SM10 λ*pir*, followed by conjugation into *P. aeruginosa* PA14 as described previously. Appropriate antibiotics were used for selection as indicated. Fungal strains outside those generated in this study were obtained from the Fungal Genetics Stock Center (Manhattan, Kansas, USA).

### *Aspergillus fumigatus* genetic manipulation

*A. fumigatus* strain A1160 *Δku80 pyrG*+ was used as the genetic background for the deletion of *gliT and gliZ*. Both mutants were generated by replacing the ORF with the dominant selection marker *hygB* (hygromycin resistance). The replacement construct was generated using overlap PCR to fuse ∼1 kb upstream and ∼1 kb downstream of the open reading frame of *gliT/gliZ* to the *hygB* marker. The resulting constructs were transformed into protoplasts of each strain, and mutants were selected for on osmotically stabilized minimal medium (GMM plus 1.2 M sorbitol) containing 200 µg/ml hygromycin B (Sigma). Protoplasts were generated using lysing enzyme and transformed as previously described. Strains were single spored and checked for correct integration, or the presence of construct in the case of the ectopic reconstituted strains via PCR.

Transformation of *A. fumigatus* with *gliT*-mCherry, *dnoP*-mCherry, *gpdA*p-mCherry or *aspf2*-mCherry into the *A. fumigatus* SH1 neutral domain was carried out using a clustered regularly interspaced short palindromic repeats (CRISPR)/Cas9-mediated protocol for gene editing, as previously described (69). crRNA was purchased from IDT (Integrated DNA Technologies, Inc.). Complete guideRNAs (gRNAs) were then assembled *in vitro* using the custom designed crRNA coupled with a commercially acquired tracrRNA. The assembled gRNAs were then combined with commercially purchased Cas9 to form ribonucleoproteins for transformation, as previously. Repair templates carrying a phleomycin resistance (*bleoR*) cassette were PCR amplified to contain 40-basepair regions of microhomology on either side for homologous integration at the double strand DNA break induced by the Cas9 nuclease. Protoplast-mediated transformations were then carried out using the hygromycin repair templates and Cas-ribonucleoproteins for gene targeting. Homologous integrations were confirmed by PCR.

### GliT and DnoP Protein expression and purification

The PCR-amplified ORFs of *gliT* and *dnoP* were cloned into the expression vector pET-19m, which introduces a TEV-cleavable N-terminal hexahistidine tag onto each protein. For purification of the His_6_-tagged proteins, the cells were grown in Turbo Broth medium (Molecular Dimensions) (1 L) at 37°C with good aeration to *A*_600_ = 0.5. The temperature was then lowered to 16°C, and isopropyl 1-thio-β-D-galactopyranoside (IPTG) was added to 1 mM final concentration to induce expression of the cloned genes. The induced cultures were grown for a further 16 h and then harvested by centrifugation (6000 × *g*, 4°C, 15 min). The cell pellet was resuspended in 20 ml of buffer A (50 mM sodium phosphate, 100 mM NaCl, 10% (v/v) glycerol, pH 8.0, containing one dissolved cOmplete™ EDTA-free Protease Inhibitor cocktail tablet (Roche). The cell lysate was clarified by centrifugation (11,000 × *g*, 4 °C, 30 min), and the filtered lysate was then loaded onto a Ni-NTA spin column (Qiagen). The column was washed five times with buffer A containing 10 mM imidazole. The His_6_-tagged proteins were eluted with buffer A containing 250 mM imidazole. Protein was concentrated and buffer exchanged to the desired concentration using an Amicon® Ultra-4 Centrifugal Filter (10 kDa NMW cut-off).

For GliT-mCherry and DnoP-mCherry expression and purification from *A. fumigatus*, 100 ml (10^5^ conidia/mL) of *A. fumigatus* strains *gliT* rec and *dnoP* rec were cultured in SDB for 24 hr, followed by 10 µM gliotoxin addition for 6 hr. Mycelia was then harvested, flash frozen in liquid nitrogen, and ground to a fine powder using a pestle and mortar. Ground mycelia was resuspended in 20 ml of buffer A and transferred to bead-beating tubes. Cells were lysed by bead beating twice for 20 s, and the tubes were placed on ice for 1 min between each homogenization. The cell lysates were clarified by centrifugation (11,000 × *g*, 4 °C, 30 min), and the filtered lysates were added to an eppendorf containing 200 µl of prewashed ALFA Selector PE magnetic beads (NanoTag Biotechnologies). Samples (4 ml) were then placed on an end-over-end orbital rotator for 1 hr at 4°C, placed on a magnetic stand, and the beads were washed five times with 1 ml of buffer A, followed by elution with 200 µl buffer A + 200 µM ALFA elution peptide. Proteins were inspected by SDS-PAGE analysis for purity.

### Dithiol oxidase enzyme assay

The dithiol oxidase activity of GliT and DnoP was measured based a method described previously (70). Briefly, the free thiol groups on reduced disulfide natural products reacts with 5,5’-dithio*bis*(2-nitrobenzoic acid) (DTNB) to yield 2-nitro-5-thiobenzoate (TNB^2-^) anions. TNB^2-^ is colored and its formation can be monitored at 412 nm. The reaction mixtures (180 µl) contained buffer (20 mM Tris-HCl, pH 7.0 and 100 mM NaCl) and TCEP-reduced substrates (gliotoxin, holomycin, chetomin, chaetocin and romidepsin) at 40 µM final. The reaction was initiated by the addition of GliT or PA4170 (to a final concentration of 0.2 µM) and incubated at room temperature for 15 min. DTNB dissolved in EtOH (5 mM) was used as a color development substrate after the oxidase reaction was complete. Enzyme free controls were used to monitor background natural product reoxidation, which was not significant over the course of the assay timeframe. The A_412_ was measured in a BioTek SynergyMx.

### Gliotoxin oxidase enzyme assay using *P. aeruginosa* cell lysates

Testing the gliotoxin oxidase activity of *P. aeruginosa* cell lysates was carried out through modifications of the above dithiol oxidase enzyme assay. Briefly. *P. aeruginosa* strains were cultured in 30 ml SCFM (static, 37C) until the cultures reached log phase (optical density at 600 nm [OD_600_] = 0.3). Gliotoxin (20 µM) or MeOH (solvent control) was then added to the cultures, which were incubated for a further 6 h. Cells from the above cultures were harvested by centrifugation (8000 × *g*, 4°C, 15 min). The cell pellet was resuspended in chilled lysis buffer (Bugbuster) containing one dissolved cOmplete™ EDTA-free Protease Inhibitor cocktail tablet (Roche). The cell lysates were clarified by centrifugation (11,000 × *g*, 4 °C, 30 min), and the final cell lysate protein content was quantified by Bicinchoninic acid (BCA) assay (Pierce™ BCA Protein Assay). Cell lysates were normalized to 10 mg/ml, and the enzyme assay was carried out immediately using fresh lysates. Reaction mixtures (200 µl) contained buffer (20 mM Tris-HCl, pH 7.0 and 100 mM NaCl) and TCEP-reduced gliotoxin at 40 µM final. The reaction was initiated by the addition of *P. aeruginosa* cell lysate (to a final concentration of 50 µg per reaction) and mixtures were incubated at room temperature for 15 min. DTNB dissolved in EtOH (5 mM) was used as a color development substrate after the oxidase reaction was complete. The A_412_ was measured in a BioTek Synergy H1 Hybrid Reader.

