## Supplementary Information for "Convergent evolution in toxin detection and resistance provides evidence for conserved bacterial-fungal interactions"

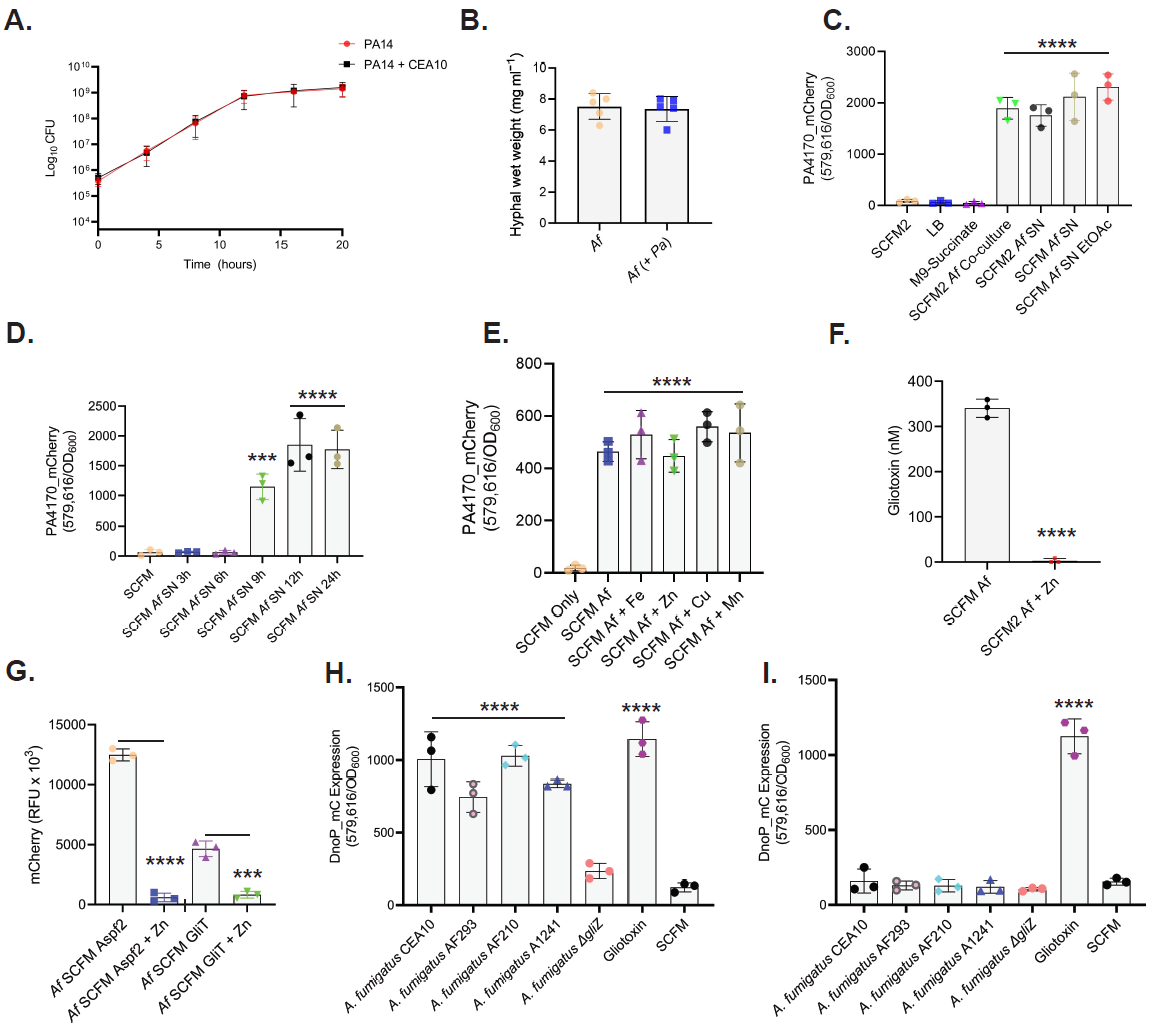

**Fig. S1. A.** CFUs of *P. aeruginosa* during growth in SCFM2 with and without the presence of *A. fumigatus*. Cultures were plated using the drop-plate method every 4 hr for 20 hr. The data are representative of three independent experiments performed in triplicate. **B.** *A. fumigatus* fungal mass (wet weight mg/ml^-1^) following 30 hr. growth in SCFM2, with or without *P. aeruginosa* coculture. The data are representative of three independent experiments performed with five replicates. Statistical significance was determined using a Student’s t test. **C.** mCherry-tagged PA4170 reporter assay examining PA4170 protein expression following growth in different laboratory media (SCFM, LB, M9-succinate), upon co-culture with *A. fumigatus* (SCFM2 *Pa*-*Af* co-culture) or using *A. fumigatus* spent culture supernatants (24 hr SCFM/SCFM2). Ethyl-acetate extracted *A. fumigatus* spent culture supernatants (24 hr) also induce PA4170 expression. Fluorescence was read at 8 hr and normalized to cell density (OD600). The data are representative of three independent experiments performed in triplicate. Statistical significance was calculated by one-way ANOVA with Dunnett’s multiple comparisons test using ‘SCFM2’ as the control column (**** P < 0.0001). Error bars represent standard deviation from the mean. **D.** mCherry-tagged PA4170 reporter fluorescence assay examining PA4170 protein expression following growth in spent *A. fumigatus* culture supernatants grown for 0-24 hr in SCFM. Fluorescence was read at 8 hr and normalized to cell density (OD600). The data are representative of three independent experiments performed in triplicate. Statistical significance was calculated by one-way ANOVA with Dunnett’s multiple comparisons test using ‘SCFM’ as the control column (*** P = 0.0006, **** P < 0.0001). Error bars represent standard deviation from the mean. **E.** mCherry-tagged PA4170 reporter assay examining PA4170 expression following *P. aeruginosa* growth in spent *A. fumigatus* SCFM culture supernatants (24 hr) supplemented with different metals (5 µM FeSO_4_, ZnSO_4_, CuSO_4_, MnSO_4_) immediately prior to *P. aeruginosa* addition. Fluorescence was read at 8 hr and normalized to cell density (OD600). The data are representative of three independent experiments performed in triplicate. Statistical significance was calculated by one-way ANOVA with Dunnett’s multiple comparisons test using ‘SCFM only’ as the control column (**** P < 0.0001). Error bars represent standard deviation from the mean. **F.** Gliotoxin production by *A. fumigatus* during growth in SCFM2 is repressed by zinc supplementation (10 µM ZnSO_4_). Gliotoxin was quantified in culture supernatants by LC-MS following 24 hr of *A. fumigatus* growth. The experiment was performed in triplicate. Statistical significance was determined using a Student’s t test (**** P < 0.0001). **G.** *A. fumigatus* is zinc limited during growth in SCFM2 and activates expression of the zinc starvation regulated protein Aspf2 and the gliotoxin biosynthetic enzyme GliT. Fluorescence assay examining mCherry-tagged Aspf2 and GliT protein expression during growth in SCFM with or without zinc supplementation (10 µM ZnSO_4_). Expression was normalized to mCherry expression (+/- 10 µM ZnSO_4_) under the control of the constitutive *gpdA* promoter. Fluorescence was read at 48 hr. The data are representative of three independent experiments performed in triplicate. Statistical significance was determined using a Student’s t test (Aspf2 +/- Zn, GliT +/- Zn) (*** P = 0.0007, **** P < 0.0001). **H.** mCherry-tagged DnoP reporter fluorescence assay examining DnoP protein expression following *P. aeruginosa* growth in spent *A. fumigatus* SCFM culture supernatants (24 h), gliotoxin (30 µM) or SCFM. Fluorescence was read at 8 hr and normalized to cell density (OD600). The data are representative of three independent experiments performed in triplicate. Statistical significance was calculated by one-way ANOVA with Dunnett’s multiple comparisons test using ‘SCFM’ as the control column (**** P < 0.0001). Error bars represent standard deviation from the mean. **I.** mCherry-tagged PA4170 reporter fluorescence assay examining PA4170 protein expression following growth in spent *A. fumigatus* SCFM culture supernatants (24 hr) supplemented with 5 µM ZnSO_4_ prior to fungal inoculation. Gliotoxin (30 µM) and SCFM wells were also supplemented with 5 µM ZnSO_4_. Fluorescence was read at 8 hr and normalized to cell density (OD600). The data are representative of three independent experiments performed in triplicate. Statistical significance was calculated by one-way ANOVA with Dunnett’s multiple comparisons test using ‘SCFM’ as the control column (**** P < 0.0001). Error bars represent standard deviation from the mean.

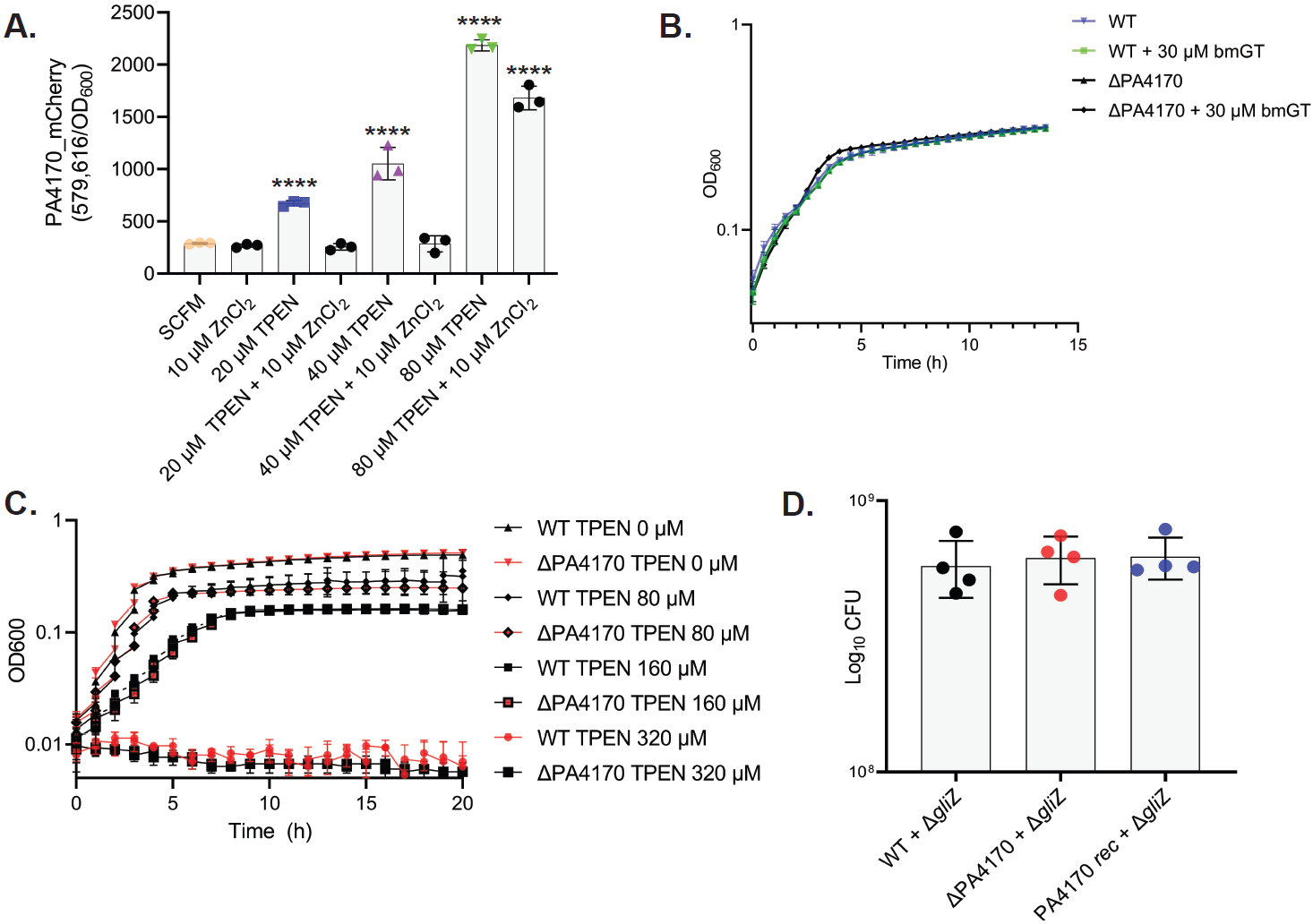

**Fig. S2. A.** Zinc starvation mediated by TPEN (N,N,N′,N′-tetrakis(2-pyridinylmethyl)-1,2-ethanediamine) supplementation activates *P. aeruginosa* PA4170-mCherry expression in a dose-dependent manner (20-80 µM). Supplementation with 10 µM ZnSO_4_ partially reverses PA4170 induction. The data are representative of three independent experiments performed in triplicate. Statistical significance was calculated by one-way ANOVA with Dunnett’s multiple comparisons test using ‘SCFM’ as the control column (**** P < 0.0001). Error bars represent standard deviation from the mean. **B.** *P. aeruginosa* wild-type or a ∆PA4170 deletion mutant are not susceptible to bisthiomethylgliotoxin exposure (30 µM). Growth curve carried out in MOPS-succinate. The data are representative of three independent experiments performed in triplicate. **C.** Deletion of PA4170 has no impact on the ability of *P. aeruginosa* to grow under TPEN mediated zinc limitation (0-320 µM). Growth curve carried out in SCFM. The data are representative of three independent experiments performed in triplicate. **D.** Growth of a ∆PA4170 deletion mutant is not impacted upon co-culture with *A. fumigatus* ∆*gliZ* when compared to wild type or PA4170 *rec* (plasmid complemented). Cultures were plated for CFU counts 20 hr after *P. aeruginosa* addition. The data are representative of three independent experiments performed in quadruplicate. Statistical significance was calculated by one-way ANOVA with Dunnett’s multiple comparisons test using ‘WT + ∆*gliZ*’ as the control column. Error bars represent standard deviation from the mean.

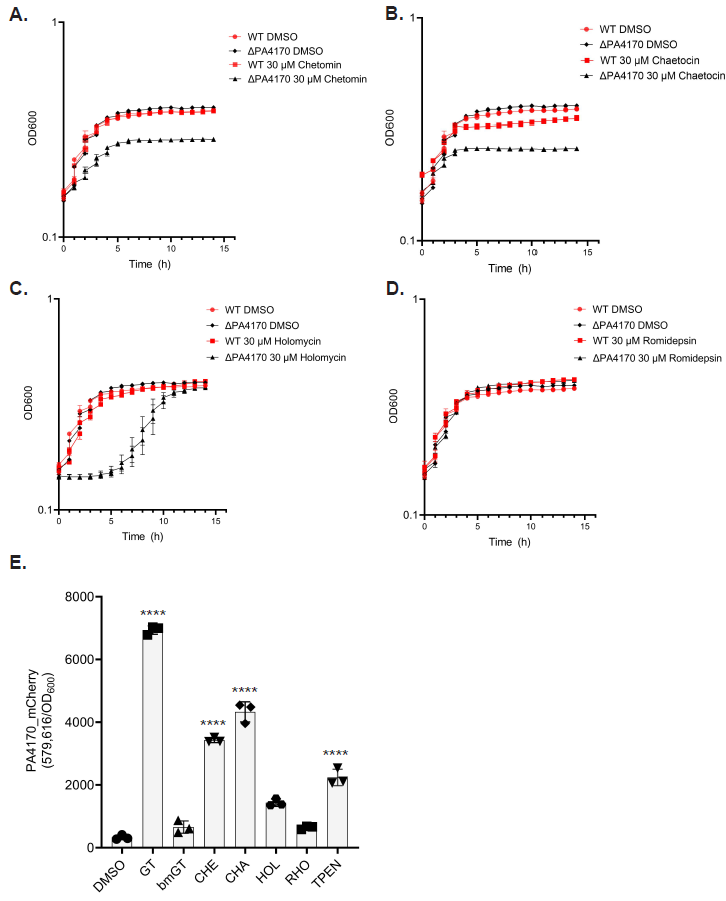

**Fig. S3. A.** Growth of a ∆PA4170 deletion mutant is more susceptible to chetomin exposure (30 µM) when compared to the *P. aeruginosa* wild type. Strains cultured in M9-succinate. The data are representative of two independent experiments performed in triplicate. **B.** Growth of a ∆PA4170 deletion mutant is more susceptible to chaetocin exposure (30 µM) when compared to the *P. aeruginosa* wild type. Strains cultured in M9-succinate. The data are representative of two independent experiments performed in triplicate. **C.** Growth of a ∆PA4170 deletion mutant is more susceptible to holomycin exposure (30 µM) when compared to the *P. aeruginosa* wild type. Strains cultured in M9-succinate. The data are representative of two independent experiments performed in triplicate. **D.** Romidepsin exposure (30 µM) has no impact on the growth of *P. aeruginosa* wild type or ∆PA4170. Strains cultured in M9-succinate. The data are representative of two independent experiments performed in triplicate. **E**. mCherry-tagged PA4170 reporter assay examining the impact of zinc supplementation (10 µM ZnSO_4_) on the ability of the ETPs gliotoxin (GT), bisthiomethylgliotoxin (bmGT), chetomin (CHE), chaetocin (CHA), holomycin (HOL) or romidepsin (RHO) to induce PA4170-mCherry expression (all ETPs added at 30 µM). TPEN was added at 50 µM. Fluorescence was read at 8 hr and normalized to cell density (OD600). The data are representative of two independent experiments performed in triplicate. Statistical significance was calculated by one-way ANOVA with Dunnett’s multiple comparisons test using ‘DMSO’ as the control column (**** P < 0.0001). Error bars represent standard deviation from the mean.

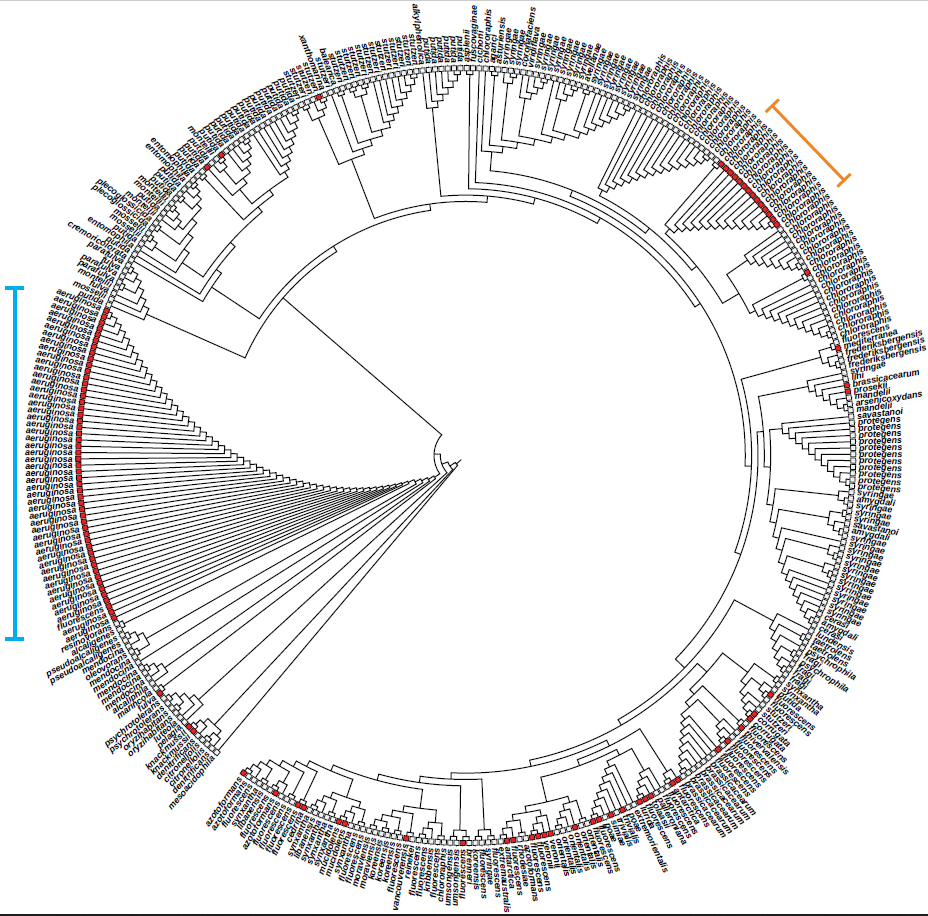

**Fig. S4.** PA4170 orthologs (red boxes) are found in most sequenced *P. aeruginosa* isolates (279/284) and are interspersed amongst other sequenced *Pseudomonas* strains (n = 74), including *P. fluorescens*, *P. antarctica*, *P. tolaasii* and *P. corrugata*. A previously unidentified phylogenetic sub-branch of *P.* *chlororaphis* which encodes a PA4170 ortholog is also indicated (orange). Maximum-likelihood tree was assembled with 16s rRNA sequences from representative, sequenced strains across the *Pseudomonas* genus.

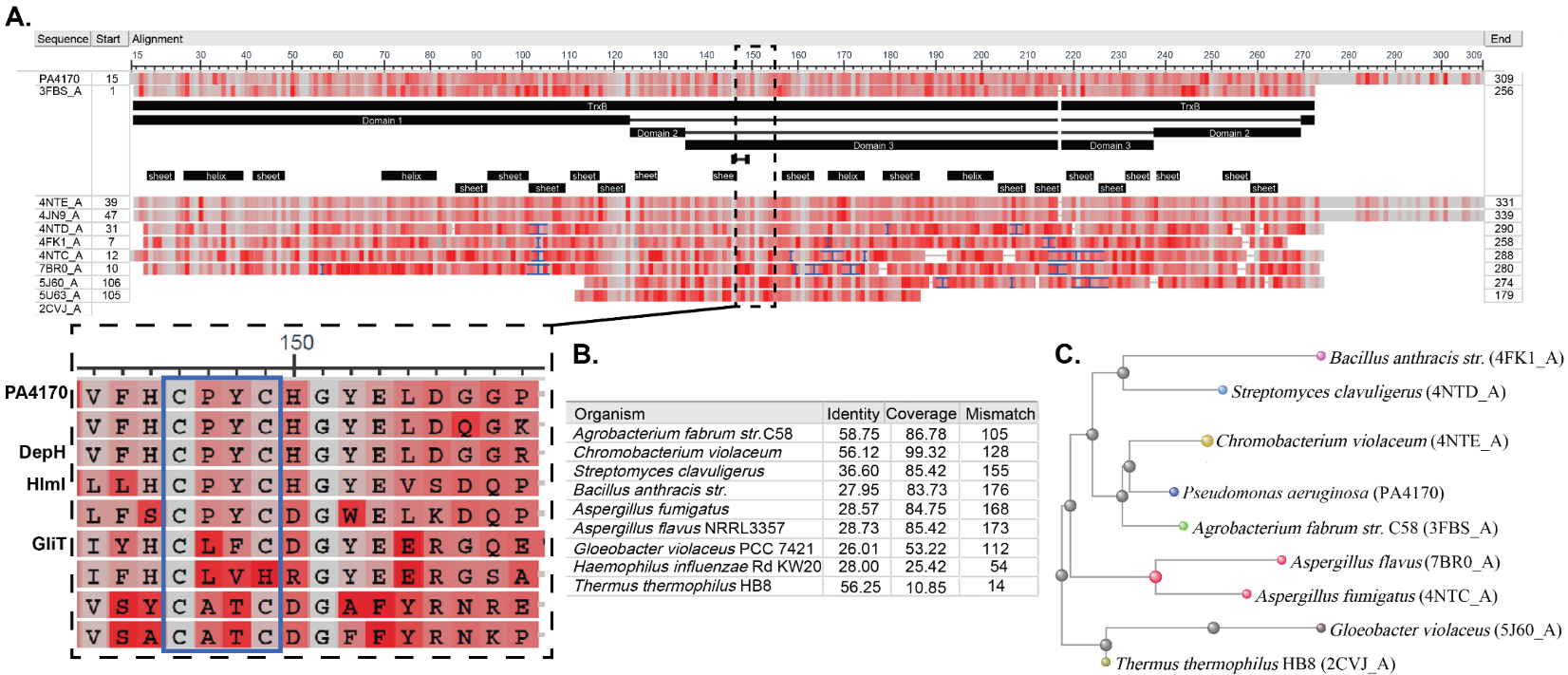

**Fig. S5. A.** BLAST search using PA4170 as a query sequence against the Protein Data Bank (PDB) identifies structurally related orthologs of PA4170 in bacteria and fungi– disulfide natural product oxidases (n = 9). Alignments wee visualized with the NCBI Multiple Sequence Alignment Viewer (MSA). The CPYC motif, characteristic of dithiol oxidases is highlighted. **B.** Sequence identity, coverage, and mismatch of the nine BLAST search PDB matches to PA4170. **C.** Phylogenetic tree of PA4170 matches as illustrated by BLAST pairwise alignment tree viewer (Fast Minimum Evolution).

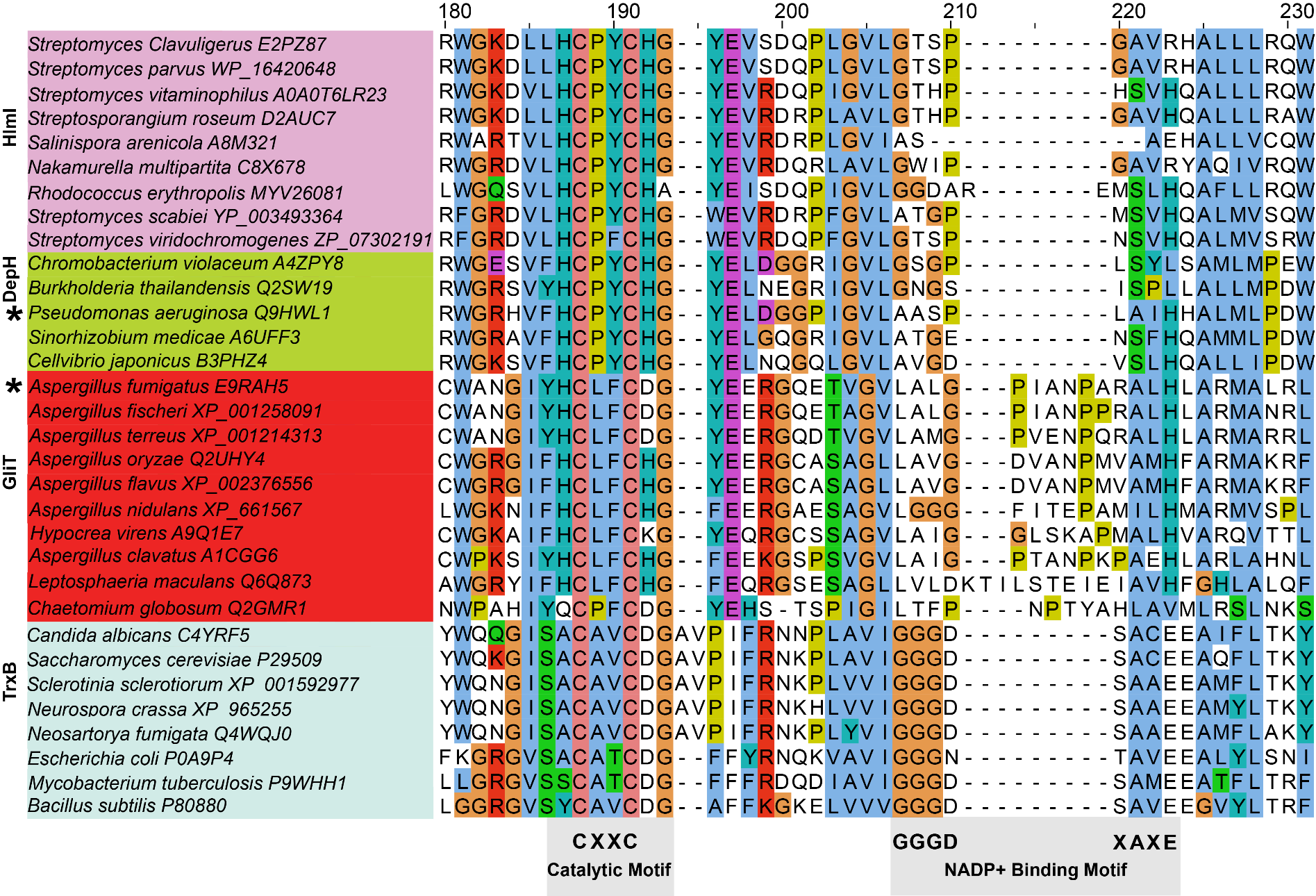

**Fig. S6.** ClustalW alignment of the catalytic residues in selected bacterial/fungal dithiol oxidases and thioredoxin reductases. The conserved CXXC catalytic motif is indicated, as is the GGGD motif specific for NADP binding in thioredoxin reductases (53). GliT (E9RAH5) and PA4170 (Q9HWL1) are highlighted with asterisks.

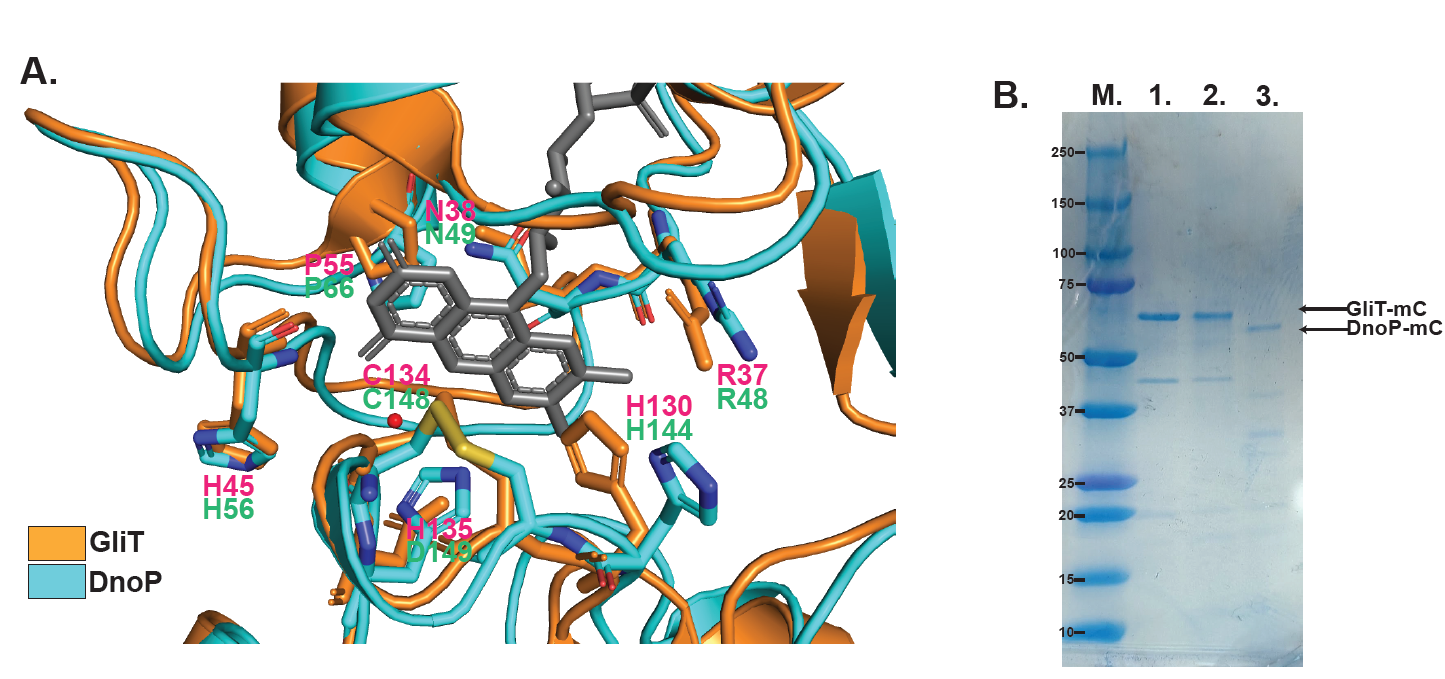
**Fig. S7. A.** GliT (4NTC) and DnoP (AlphaFold) structure overlay (PyMOL) demonstrating the positions of key shared active site residues (R37/R48, H130/H144, H45/H56, P55/P66, N38/N49, C134/C148) and alterations (H135/D149). FAD shown in grey. **B.** SDS-PAGE gel demonstrating successful ALFA-tag mediated purification of active GliT-mCherry and DnoP-mCherry from *A. fumigatus* strains *gliT* rec and *dnoP* rec, respectively. Lane M; molecular marker. 1; GliT-mCherry A (≈63 kDa), 2; GliT-mCherry B (≈63 kDa), 3; DnoP-mCherry (≈55 kDa).

**Table S1.** Strains and plasmids used in this study.

| **Strain name** | **Description** | **Source** |
| --- | --- | --- |
| DH5α | *E. coli* cloning strain | Zymo |
| BL21(DE3) | *E. coli* protein expression strains | NEB |
| SM10 (λ pir+) | *E. coli* donor strain for conjugation | (1) |
| *P. aeruginosa* PA14 | Wild-type laboratory strain | (2) |
| PA14 Δ*PA4170* | *PA4170* insertion mutant in PA14 | This study |
| PA14 Δ*PA4170 :: PA4170* | PA14 Δ*PA4170* carrying plasmid pUCP20 with PA4170. *amp^R^* | This study |
| *P. aeruginosa* PAO1 | Wild-type laboratory strain | (3) |
| *P. aeruginosa* C5912M | Clinical isolate from CF patient (sputum, mucoid) | (4) |
| *P. aeruginosa* C0324C | Clinical isolate from CF patient (sputum, classic) | (4) |
| *PA14 PA4170-mC* | *PA4170-ALFA-*mCherry translational fusion in PA14 | This study |
| FGSC A1260 (CEA10) | *A. fumigatus* Wild type: MAT1-1 (also known as CEA10, CBS 144–89CBS 144.89, AF10). Isolated from patient with IPA. | (5) |
| FGSC A1241 | *A. fumigatus* wild type AfIR964 | FGSC |
| FGSC A1259 | *A. fumigatus*, clinical isolate AF210 (NCPF 7101) | FGSC |
| FGSC A1100 | *A. fumigatus* wild type AF293 | FGSC |
| A1160 Δ*ku80* *pyrG*+ | *A. fumigatus* strain derived from A1260 that lacks nonhomologous end joining (*Δku80*). Δ*akuBku80*::*pyrG*−-*zeo*, *pyrG*−::*pyrGAf*; *MAT1-1,* MFIG001 | (6) |
| A1160 ∆*gliT*::*hygB* | A1160 *gliT* deletion strain, *hygB* | This study |
| A1160 ∆*gliT*::*hygB; gliT-mC::bleoR* | A1160 *gliT* deletion strain complemented with *gliT*-*mCherry* (SH1 region), *hygB*:*bleoR* | This study |
| A1160 ∆*gliT*::*hygB; gliTp-PA4170-mC::bleoR* | A1160 *gliT* deletion strain in A1160 with *gliTp-PA4170*-*mCherry* inserted (SH1 region), *hygB*:*bleoR* | This study |
| A1160 ∆*gliZ*::*hygB* | A1160 *gliZ* deletion strain, *hygB* | This study |
| A1160 *aspf2*-mC*::bleoR* | A1160 with *aspf2*-*mCherry* inserted in the CEA10 SH1 region, *bleoR* | This study |
| A1160 *gpdAp*-mC*::bleoR* | A1160 with *gpdAp*-*mCherry* inserted in the CEA10 SH1 region, *bleoR* | This study |
| **Plasmid name** | **Description** | **Source** |
| pUCP20 | *E. coli* – *Pseudomonas* shuttle vector for constitutive expression of cloned genes; Amp^R^ | (7) |
| pSKD1 | PA4170 promoter and ORF cloned into pUCP20; Amp^R^ | This study |
| pET19m | Modified pET19 (Invitrogen), containing TEV protease-cleavable N terminally His6-tag; Amp^R^ | (8) |
| pSKD2 | *A. fumigatus* *gliT* ORF cloned into pET-19m; Amp^R^ | This study |
| pSKD3 | *P. aeruginosa* *PA4170* ORF cloned into pET-19m; Amp^R^ | This study |
| pEXG2 | Parental plasmid used markerless DNA mutagenesis; *sacB*, Gen^r^ | (9) |
| pSKD4 | pEXG2-Δ*PA4170*; *PA4170* deletion cassette; *sacB*, Gen^r^ | This study |
| pSKD5 | pEXG2-*PA4170-mC; PA4170* mCherry translational fusion cassette; *sacB*, Gen^r^ | This study |
| pMF440 | Broad host range plasmid for constitutive expression of mCherry; Amp^R^ | (10) |
| pAN 7.1 | Hygromycin resistant cassette source for PCR amplification; Amp^R^ | (11) |
| pAN 8.1 | Phleomycin resistant cassette source for PCR amplification; Amp^R^ | (12) |
| pSKD6 | *gliT* promoter and *gliT* ORF fused with an ALFA tag linker and C-terminus mCherry. Plasmid contains *bleoR* (from pAN 8.1); Amp^R^ | This study |
| pSKD7 | *gliT* promoter and *PA4170* ORF fused with an ALFA tag linker and C-terminus mCherry. Plasmid contains *bleoR* (from pAN 8.1); Amp^R^ | This study |
| pSKD8 | *aspf2* promoter and *aspf2* ORF fused with an ALFA tag linker and C-terminus mCherry. Plasmid contains *bleoR (*from pAN 8.1); Amp^R^ | This study |
